# Resting-state “Physiological Networks”

**DOI:** 10.1101/660787

**Authors:** Jingyuan E. Chen, Laura D. Lewis, Catie Chang, Nina E. Fultz, Ned A. Ohringer, Bruce R. Rosen, Jonathan R. Polimeni

**Affiliations:** Athinoula A. Martinos Center for Biomedical Imaging, Massachusetts General Hospital, Boston, MA, USA; Department of Radiology, Harvard Medical School, Boston, MA, USA; Department of Biomedical Engineering, Boston University, Boston, MA, USA; Department of Electrical Engineering and Computer Science, Vanderbilt University, Nashville, TN, USA; Harvard-Massachusetts Institute of Technology Division of Health Sciences and Technology, Cambridge, MA, USA

**Keywords:** heart rate, global signal regression, fMRI, resting state functional connectivity, respiratory variation

## Abstract

Slow changes in systemic brain physiology can elicit large fluctuations in fMRI time series, which may manifest as structured spatial patterns of temporal correlations between distant brain regions. These correlations can appear similar to large-scale networks typically attributed to coupled neuronal activity. However, little effort has been devoted to a systematic investigation of such “physiological networks”—sets of segregated brain regions that exhibit similar physiological responses—and their potential influence on estimates of resting-state brain networks. Here, by analyzing a large group of subjects from the 3T Human Connectome Project database, we demonstrate brain-wide and noticeably heterogenous dynamics attributable to either respiratory variation or heart rate changes. We show that these physiologic dynamics can give rise to apparent “connectivity” patterns that resemble previously reported resting-state networks derived from fMRI data. Further, we show that this apparent “physiological connectivity” cannot be removed by the use of a single nuisance regressor for the entire brain (such as global signal regression) due to the clear regional heterogeneity of the physiological responses. Possible mechanisms causing these apparent “physiological networks”, and their broad implications for interpreting functional connectivity studies are discussed.

## 1 Introduction

During the resting state, defined experimentally as a state in which participants are not given any explicit stimulus or task, distributed brain regions exhibit temporally synchronized hemodynamic changes that can be measured by functional magnetic resonance imaging (fMRI) (Biswal B et al. 1995; Fox MD and ME Raichle 2007). Although the biophysical origins of fMRI-based functional connectivity (FC) are not completely understood, these seemingly spontaneous fluctuations in hemodynamics—and their correlations across brain regions—have been demonstrated by many studies to reflect neural activity. For instance, in animal models, the amplitudes of regional spontaneous blood oxygenation levels track the power of electrophysiological recordings across several frequency bands (e.g., (Shmuel A and DA Leopold 2008; Mateo C et al. 2017)), and large-scale correlations in the hemodynamics across brain regions closely match the correlations seen in concurrent neuronal calcium dynamics in rodents (Ma Y et al. 2016). In human studies, the existence of spontaneous, task-free FC among distributed brain regions have also been confirmed by alternative neuroimaging modalities such as electrocorticography (He BJ et al. 2008; Miller KJ et al. 2009; Kucyi A et al. 2018) and magnetoencephalography (Brookes MJ et al. 2011; Hipp JF et al. 2012; Baker AP et al. 2014). Therefore, these correlations in hemodynamic fluctuations appear to in part reflect correlations in the underlying neuronal activity.

Although this evidence points to a neural origin of some components of resting-state fMRI signal fluctuations, hemodynamic signals provide an indirect measure of neural activity and non-neuronal factors can also drive changes in blood flow and oxygenation. These factors should also be considered when interpreting resting-state FC based on fMRI. Systemic physiological changes, including those associated with cardiac and respiratory cycles, are well known to influence hemodynamic changes through multiple mechanisms. In particular, low-frequency fluctuations in end-tidal CO_2_ levels (Wise RG et al. 2004), respiration volume (Birn RM et al. 2006), and heart rate (Shmueli K et al. 2007) have been shown to account for considerable variance in BOLD fMRI signals during both the resting state as well as during tasks (see (Liu TT 2016; Caballero-Gaudes C and RC Reynolds 2017) for reviews).

Recent studies have revisited the physiological contribution to fMRI-based resting-state observations. In lieu of directly quantifying the exact BOLD variance explained by different physiological elements, these studies focused on the spatial heterogeneity of BOLD responses linked to different physiological dynamics, and have identified structured, regular spatial patterns related to physiology. For example, by applying independent component analysis (ICA) to synthesized datasets simulating the propagation of total hemoglobin (deoxyhemoglobin plus oxyhemoglobin) across the entire brain, Tong et al. identified several spatial components that closely resembled common resting-state networks (RSNs), such as the default-mode network (DMN) and visual network (Tong Y et al. 2015). By introducing a minor hypercapnic challenge with a paradigm orthogonal to concurrent cognitive stimuli, Bright et al. identified spatial patterns of the BOLD response that overlapped with task-(de)activated neuronal networks but could be ascribed specifically to physiological responses (Bright MGM, K. 2014).

The rationale behind these studies is that vascular responses following systemic physiological changes are not spatially homogeneous (Chang C et al. 2009; Pinto J et al. 2017), by virtue of both inconsistent vascular arrival time (MacIntosh BJ, N Filippini, et al. 2010; Tong Y et al. 2017) and regional differences in the temporal shape/delay of the BOLD responses that may be determined in part by local vascular anatomy and density (Aguirre GK et al. 1998; Handwerker DA et al. 2004). Yet, common approaches employed to derive functional networks from BOLD fMRI data (i.e., either seed-based linear Pearson correlation analyses or data-driven independent component analyses) cannot discern “apparent” temporal synchrony caused by consistent vascular responses from the targeted synchrony caused by coupled neuronal activity. For example, if two distinct brain regions happen to exhibit similar BOLD responses to heart rate changes, which would result in coherent cardiac-driven BOLD fluctuations between these two regions, even in the absence of any direct or indirect coupling of the neuronal activity between these regions, it is likely that these two regions would appear to be functionally connected when using conventional FC estimation approaches. Therefore, systemic physiological changes alone can potentially yield network-like spatial patterns depending on the spatial pattern of the physiological responses seen in the BOLD data (Tong Y *et al*. 2015). In the remainder of this article, we therefore term such apparent temporal synchrony linked with systemic physiological fluctuations as “physiological connectivity”, to differentiate it from the conventional notion of purely neuronally-driven fluctuations that give rise to networks of functionally connected brain regions. Note that we do not preclude potential neural information carried by such “physiological” dynamics, and indeed, neuronal controls involved in regulating brain physiology (reflecting, e.g., autonomic outflow (Cechetto DFSe, C.B. 1990; Beissner F et al. 2013)) will elicit hemodynamic changes that can have mixtures of neuronal and physiological origins.

Today, a comprehensive characterization of apparent “network” structures that arise from specific physiological or non-neuronal processes is lacking. Characterizing these apparent network structures with a putative physiological origin, however, may be of critical importance for the successful identification of meaningful functional connectivity-based measures of brain function, particularly for neurological conditions or patient populations associated with altered or atypical brain physiology (e.g., subjects with varying vigilance (Oken BS et al. 2006; Chang C et al. 2018), stress levels (Wang J et al. 2005), or with cardiovascular diseases (MacIntosh BJ et al. 2010; Lv Y et al. 2013; Al-Bachari S et al. 2014; Christen T et al. 2015; Jahanian H et al. 2018)).

The primary goal of the present work is to offer a systemic examination of the spatiotemporal dynamics in BOLD fMRI associated with changes in two specific “physiological” signals—respiratory volume and heart rate. We analyzed a large group of subjects from the 3T WU-UMinn *Human Connectome Project* (HCP) dataset (Van Essen DC et al. 2013). With the characterized “physiological” dynamics, we identified a detailed topology of potential network-like patterns that may arise in the absence of overtly neutrally-driven processes; and showed that these “physiological networks” resemble common resting-state networks (RSNs) to some extent. We also caution that global signal regression (GSR), a controversial but still widely-used means to remove artifacts present globally in the resting-state fMRI data, can amplify spatial heterogeneity in these patterns, and is therefore not sufficient to eliminate the influence exerted by “physiological” dynamics.

## 2 Materials and Methods

### 2.1 Data and pre-processing

Resting-state fMRI data of 190 subjects from the 3T WU-UMinn HCP were examined in this study (Smith SM et al. 2013; Van Essen DC et al. 2013). A list of employed subject ID numbers is provided in Supplementary Table S1. Each subject was scanned during two experimental sessions; here we analyzed one 15-min resting-state BOLD fMRI scan of the first session (during which subjects were instructed to keep their eyes open and fixate on cross-hair fixation target). Functional MRI data were collected using a gradient-echo Simultaneous Multi-Slice EPI sequence with the following parameters: 2 mm isotropic resolution, TR = 0.72 s, TE = 33.1 ms, multi-band factor = 8, flip angle = 52°, 72 slices, echo spacing 0.58 ms, and a left-to-right phase encoding direction. Cardiac signals were measured by a pulse oximeter placed on the fingertip and respiratory signals were measured by a bellow placed around the chest, with 400 Hz sampling rates for both recordings. All downloaded data had been processed with the “minimal preprocessing pipeline” of the HCP (Glasser MF et al. 2013) and had not undergone any physiological denoising.

### 2.2 Identifying the spatiotemporal BOLD dynamics coupled with slow physiological changes

#### 2.2.1 Extracting respiratory variation (RV) and heart-beat-interval (HBI) signals from sensor recordings

The temporal standard deviation of respiratory waveforms (from the chest bellows traces) and the averaged inter-heart beat intervals (from the pulse oximetry traces) were computed within a sliding window of 6 s centered at the time of each TR (i.e., every 0.72 s) to yield the time-varying estimates of RV and HBI, as illustrated in Fig. 1A. Although both respiratory and cardiac cycles occur at frequencies well above 0.1 Hz, RV and HBI fluctuations vary primarily at much slower temporal scales that overlap with spontaneous fluctuations, as shown in Fig. 1B.

**Figure 1:**
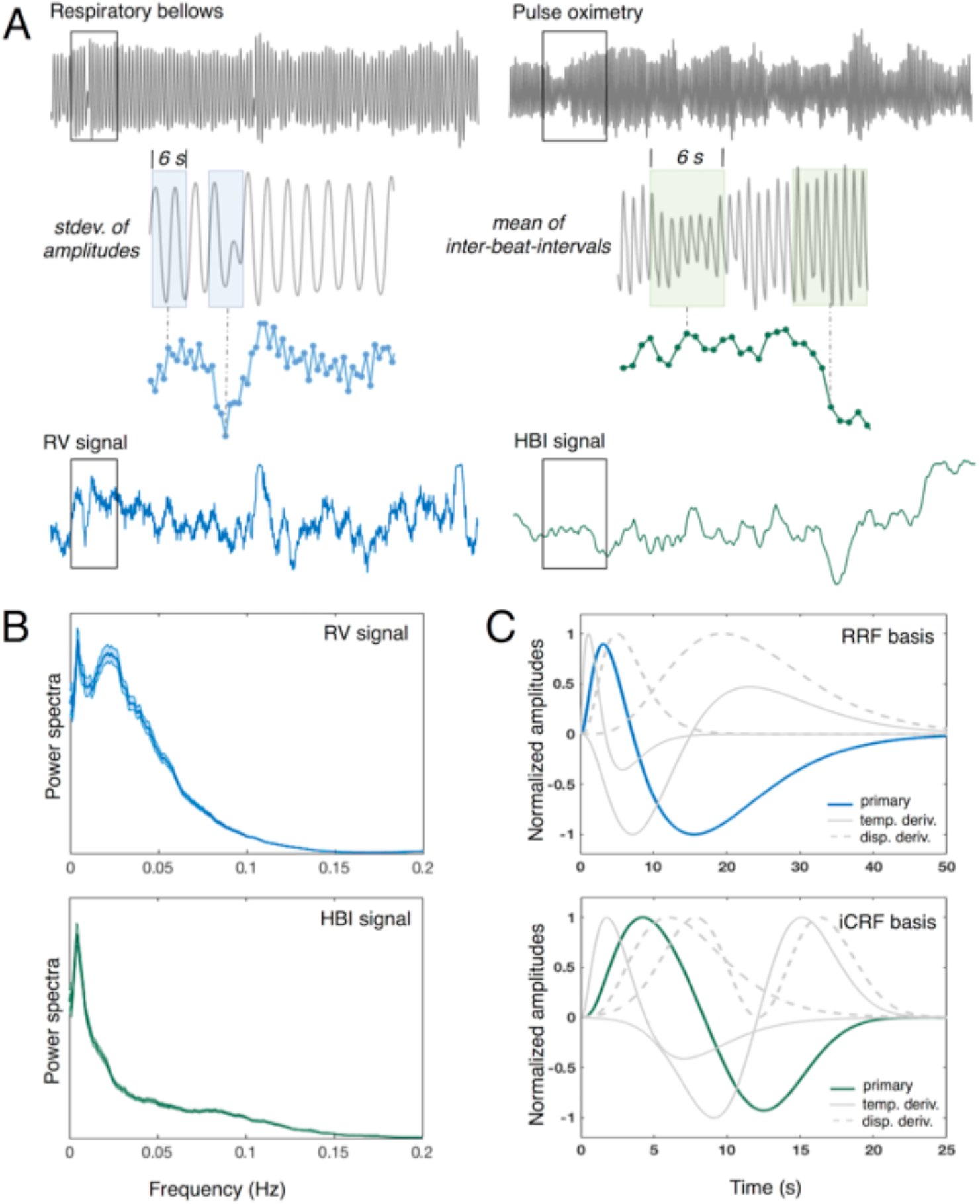
(A) Illustration of the extraction of RV and HBI time courses from recorded respiration and pulse waveforms (in one exemplar subject). (B) Mean and standard errors of the RV and HBI power spectra across 190 subjects. (C) Basis functions used to fit voxel-wise RRFs and iCRFs, including the primary basis element (i.e., the standard RRF and CRF reported previously, blue or green solid line), the temporal derivative (gray solid line) and the dispersive derivative (gray dashed line).

#### 2.2.2 Characterizing voxel-wise BOLD responses elicited by RV and HBI changes

Impulse responses following RV and HBI changes, termed as the respiratory response function (RRF) and inverse cardiac response function (iCRF, given that HBI is anti-correlated with heart rates), were jointly derived using ordinary least squares fitting-based linear regression, with basis functions shown in Fig. 1C and supplementary Eqn. S1. This basis set was generated from the standard RRF and CRF reported previously (Birn RM et al. 2008; Chang C *et al*. 2009) combined with additional temporal and dispersive derivatives generated analytically from the reported formula for each response function (each consisting of a weighted sum of two Gamma, or one Gamma and one Gaussian functions). Quasi-periodical physiological fluctuations time-locked to respiratory and cardiac cycles were modeled by RETROICOR (Glover GH et al. 2000), and were included, along with temporal drifts, as additional covariates in the regression model. To achieve comparable results across subjects, RV and HBI waveforms were standardized to Z-scores (i.e., de-meaned and divided by the standard deviation), and the BOLD time-series data were normalized to percent signal changes prior to the linear regression analyses. Voxel-wise RRFs and iCRFs were evaluated for data both with and without spatial smoothing (isotropic 3D Gaussian kernel, FWHM = 5 mm). To generate a null condition of RRF and iCRF shapes characterized by the proposed approach, as a control analysis the same de-convolution procedure was repeated in each subject’s dataset with RV and HBI measures from a different subject from the same cohort.

### 2.3 Assessing structured spatial patterns of RRFs and iCRFs

Having obtained the spatiotemporal patterns linked with RV and HBI changes, i.e., the derived voxel-wise RRFs and iCRFs, we next explored whether these “physiologically”-driven fluctuations in the BOLD fMRI data possibly generate apparent networks—defined as multiple sets of distributed brain regions that exhibit similar responses to RV or HBI changes. K-means spatial clustering was performed on the group-mean voxel-wise RRFs and iCRFs, with cluster number *k* varied from 2 to 10 to assess any hierarchical structures present within the “physiological” dynamics.

### 2.4 Comparing apparent connectivity driven by RV changes with functional connectivity associated with neuronal activity

Analyses described in this section were motivated by the observation (see Section 3.2 below) that two prominent and segregated cortical clusters emerge from the RRF dynamics, with one consisting of primary sensory cortical areas and the other consisting of cortical areas involved in higher cognitive functions. We therefore performed a more in-depth examination of the apparent connectivity patterns arising from the slow respiratory dynamics, and compared them with the functional networks estimated from spontaneous neuronal activity.

#### 2.4.1 Constructing “purely” RV-related dynamics

To simulate BOLD fMRI data containing only slow respiratory fluctuations similar to those observed in our measured resting-state data, we convolved the *group-average* voxel-wise RRFs with the recorded RV waveforms of each HCP subject, resulting in 190 synthesized fMRI datasets containing purely RV-related dynamics (and no additional fluctuations such as those driven by spontaneous neuronal dynamics or other physiological dynamics such as the cardiac cycle).

#### 2.4.2 Constructing “clean” spontaneous neuronal activity

In addition to the minimal preprocessing pipeline, BOLD data for each HCP subject were further de-noised by linearly projecting out temporal drifts, RETROICOR co-variates, and RV/HBI-related slow physiological fluctuations, i.e., de-noised data consisting of the residuals of the linear regression analysis described above in Section 2.2.2 were considered as our best estimate of “clean” BOLD fluctuations mainly reflecting spontaneous neuronal activity.

#### 2.4.3 Comparing connectivity patterns of RV-related dynamics and spontaneous fluctuations

Having generated time-series data containing either “purely” RV-related dynamics or “clean” spontaneous fluctuations, we next computed both seed-based and network-based connectivity caused by either source of resting-state BOLD signal changes.

Linear Pearson correlation with respect to two seed regions—one centered within the primary visual cortex (6-mm-radius sphere centered at MNI (−2, −72, 10)), and one centered within dorsal posterior cingulate cortex (6-mm-radius sphere centered at MNI (0, −60, 46) (Chen JE et al. 2017))—was calculated for both the “purely” RV-related dynamics and “clean” spontaneous fluctuations datasets across all brain voxels for each subject. These seed locations were chosen such that both connectivity patterns of the two major RV-related clusters identified in our resting-state BOLD data (see Section 3.2 below) would be revealed.

To assess the overall inter-network connectivity patterns generated by either source of resting-state BOLD signal changes, we first parcellated the cerebral cortex into 17 networks in MNI152 volume space using the Yeo et al. 2011 RSN atlas reported previously (Yeo BT et al. 2011; Buckner RL et al. 2011), then evaluated the connectivity matrix amongst the sets of representative network signals within both the “purely” RV-related dynamics and “clean” spontaneous fluctuations datasets derived by averaging the time series across all voxels contained within each network and then computing pairwise correlations of the averaged signals from all 17 networks.

Both the subject-level seed-based and the subject-level network-based correlation estimates were Fisher Z-transformed, then both were entered into a random-effects t-test and also averaged across all the subjects, yielding group-level estimates of connectivity driven by RV-related dynamics and by spontaneous fluctuations.

To test whether simple nuisance regression using a single waveform for the entire brain can remove the apparent connectivity related to systemic physiological fluctuations, identical analyses were then performed for the “purely” RV-related dynamics dataset and, as a reference, for “clean” spontaneous fluctuations dataset, after regressing out the average signal of all brain voxels (i.e., global signal regression (GSR)). This additional analysis was motivated by the common assumption that GSR can minimize BOLD fluctuations triggered by global nuisance factors, such as systemic physiological changes. However, given the inconsistent spatial patterns of RRFs across the brain, we expected that GSR may not be effective at eliminating regional respiratory fluctuations, and that when applied to synthetic data consisting of “purely” RV-related dynamics, the residual fluctuations would simply show an altered correlation pattern that can still resemble large-scale networks expected from synchronous neuronal activity.

## 3 Results

### 3.1 Spatiotemporal BOLD dynamics coupled with RV and HBI changes

Group-averaged spatiotemporal patterns of RV- and HBI-related brain dynamics are shown in Fig. 2A (with spatial smoothing) and Supplementary Fig. S1 (without spatial smoothing). Although the RRFs and iCRFs both relate to systemic physiological changes, both response functions exhibit clear heterogeneous amplitudes and delays across the brain. Following a positive change in RV, most brain regions underwent an initial signal increase and a subsequent undershoot that slowly recovered to the baseline condition. Notably, the temporal evolution of the respiratory response within the brainstem, thalamus, anterior cingulate cortex and primary sensory regions preceded the response seen in frontoparietal cortex and the ventricles by several seconds (Fig. 2A, top). In contrast, BOLD changes accompanying an HBI increase (i.e., a heart rate decrease) were more consistent across the cerebral cortex—most regions exhibited a bimodal pattern with an initial negative response, and a subsequent positive response. The cardiac responses within the large blood vessels, white matter and ventricles exhibited an approximately opposite or delayed trend (Fig. 2A, bottom). The heterogeneous temporal profiles of the voxel-wise RRFs or iCRFs are summarized in Fig. 2B.

**Figure 2:**
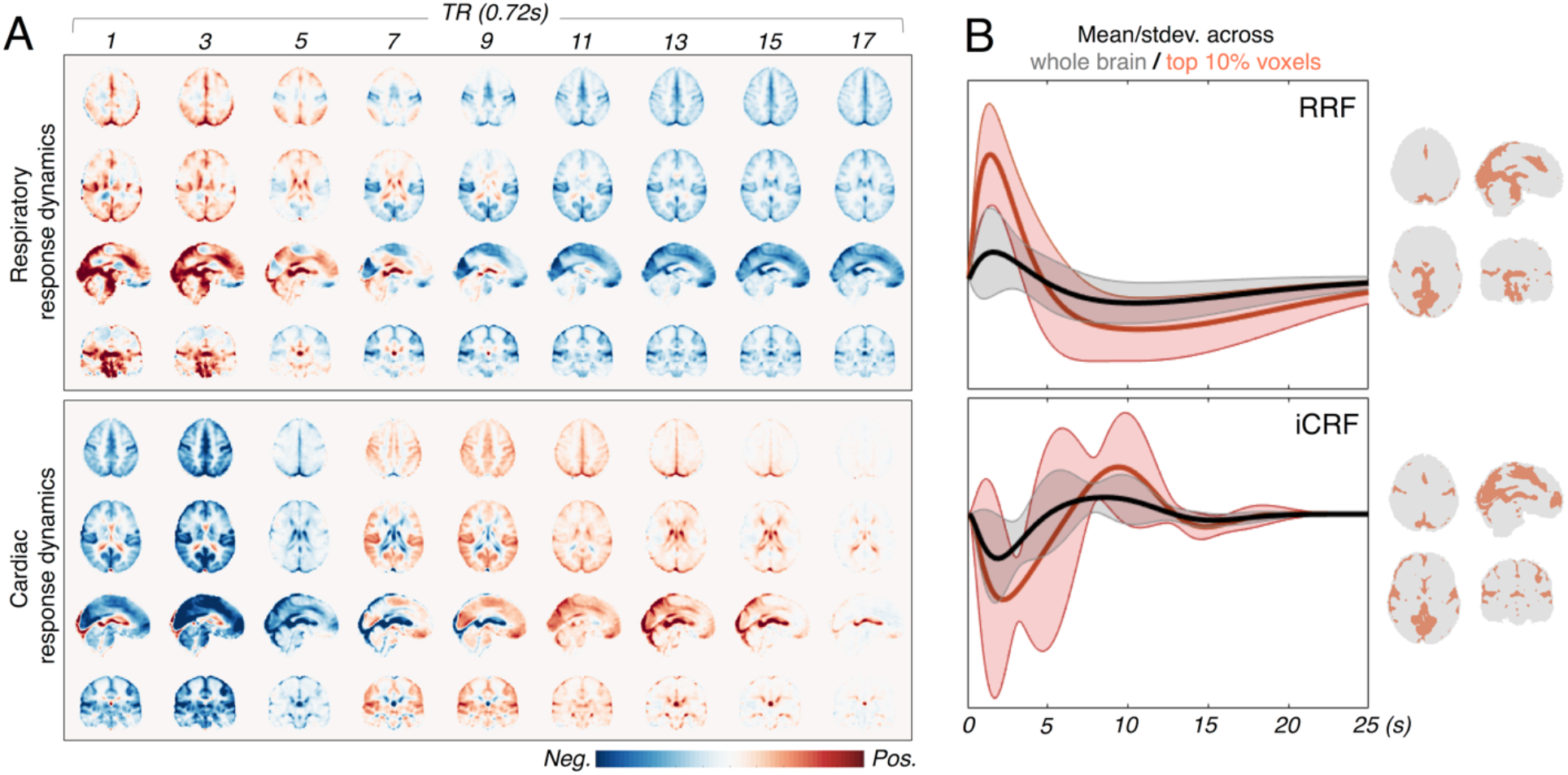
Spatially structured fMRI signals following RV and HBI changes. (A) Spatiotemporal dynamics of voxel-wise RRFs and iCRFs averaged across 190 subjects (after smoothing with an isotropic 3D Gaussian kernel, FWHM = 5 mm). (B) Mean and standard deviation of the RRFs and iCRFs across the whole brain (gray) and across voxels whose maximum/minimum (of RRFs/iCRFs respectively) values are amongst the top 10% (red).

As a control analysis, responses were measured in each subject based on the RV or HBI curves derived from a different subject, and these responses exhibited minimal levels of signal changes (an exemplar case is shown in Supplementary Fig. S2). This indicates that the observed response functions derived from the BOLD data and the physiological recordings measured in the same subject, shown in Fig. 2, are indeed related to the RV and HBI dynamics.

### 3.2 RRF and iCRF “networks”

Given the salient respiratory- and cardiac-associated dynamics we observed in the fMRI signals, we next asked how these patterns were spatially organized, using spatial clustering to identify large-scale patterns. RRF and iCRF clusters derived from K-means clustering are shown in Fig. 3. Results for *k* = 2 clusters reflect the pattern of the responses that can be seen by inspecting the spatial patterns of the responses presented in Fig. 2A: the cortical RRFs were separated into primary sensory regions (‘red’) and frontoparietal regions commonly engaged in higher cognitive functions (‘blue’); whereas the iCRF dynamics were divided into cortical regions (‘red’) and the rest of the brain (‘blue’). It is also noteworthy that higher values of *k* continue to parcellate the cortex into separate components with distinguishable waveforms for both physiological dynamics, and that the voxels within each cluster tend to organize in ways that can resemble classic FC networks (*k* = 4, 10).

**Figure 3:**
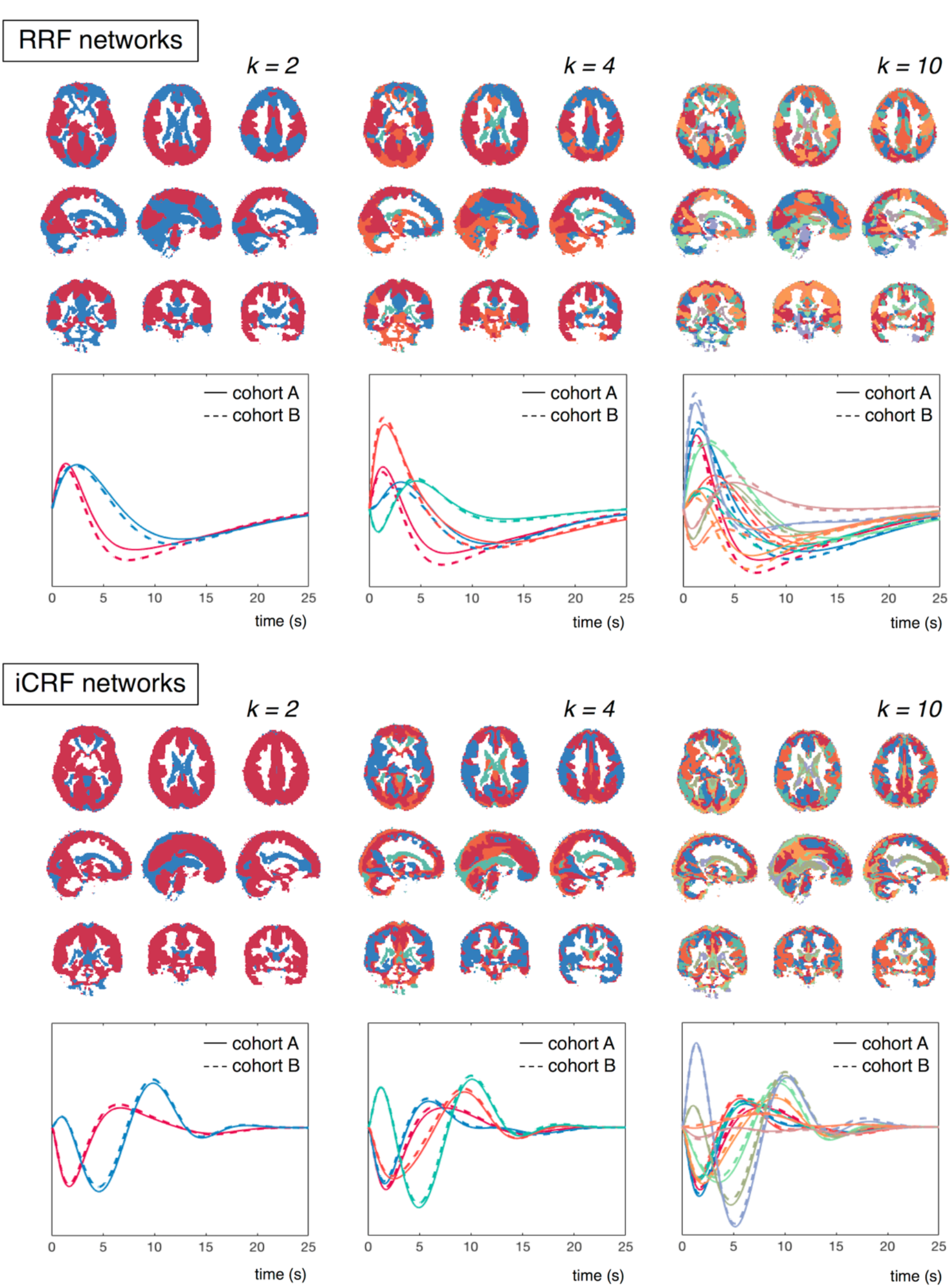
RRF and iCRF networks identified through K-means clustering using different numbers of clusters (*k* = 2, 4, 10). Cluster membership for each voxel is indicated by a color, with each cluster assigned a distinct color, and the mean waveform within each cluster is colored accordingly below the spatial map (cohort A: results from the first cohort of 95 subjects; cohort B: results from the second cohort of 95 subjects).

To validate the reliability of the derived RRFs and iCRFs for each cluster, we additionally separated the 190 HCP subjects into two cohorts with equal numbers of participants. The physiological responses averaged within each spatial cluster of both cohorts are shown in Fig. 3; both the RRFs and iCRFs were remarkably similar between the two cohorts (‘cohort A’ vs. ‘cohort B’), indicating the consistency of the observed spatial segregation of RRF/iCRF patterns.

### 3.3 Apparent connectivity drive by RV-related dynamics resembles functional connectivity in spontaneous fluctuations

To determine how these physiological fluctuations might influence apparent FC measures, we calculated seed-based connectivity patterns and network-based connectivity matrices, derived from both the synthetic data containing “purely” RV-related dynamics and the measured data containing primarily “clean” spontaneous fluctuations (see section 2.4 above). The results are shown in Fig. 4 and Fig. 5. Although RV-related dynamics were simulated with recorded RV changes instead of temporally white random processes (and therefore more weighted toward low-frequency fluctuations), the resulting connectivity patterns were still very consistent with the clustering results of voxel-wise RRFs as can be seen by comparing the RV dynamics networks (without GSR) shown in Fig. 4 with the *k* = 2 RRF network clusters shown in Fig. 3.

**Figure 4:**
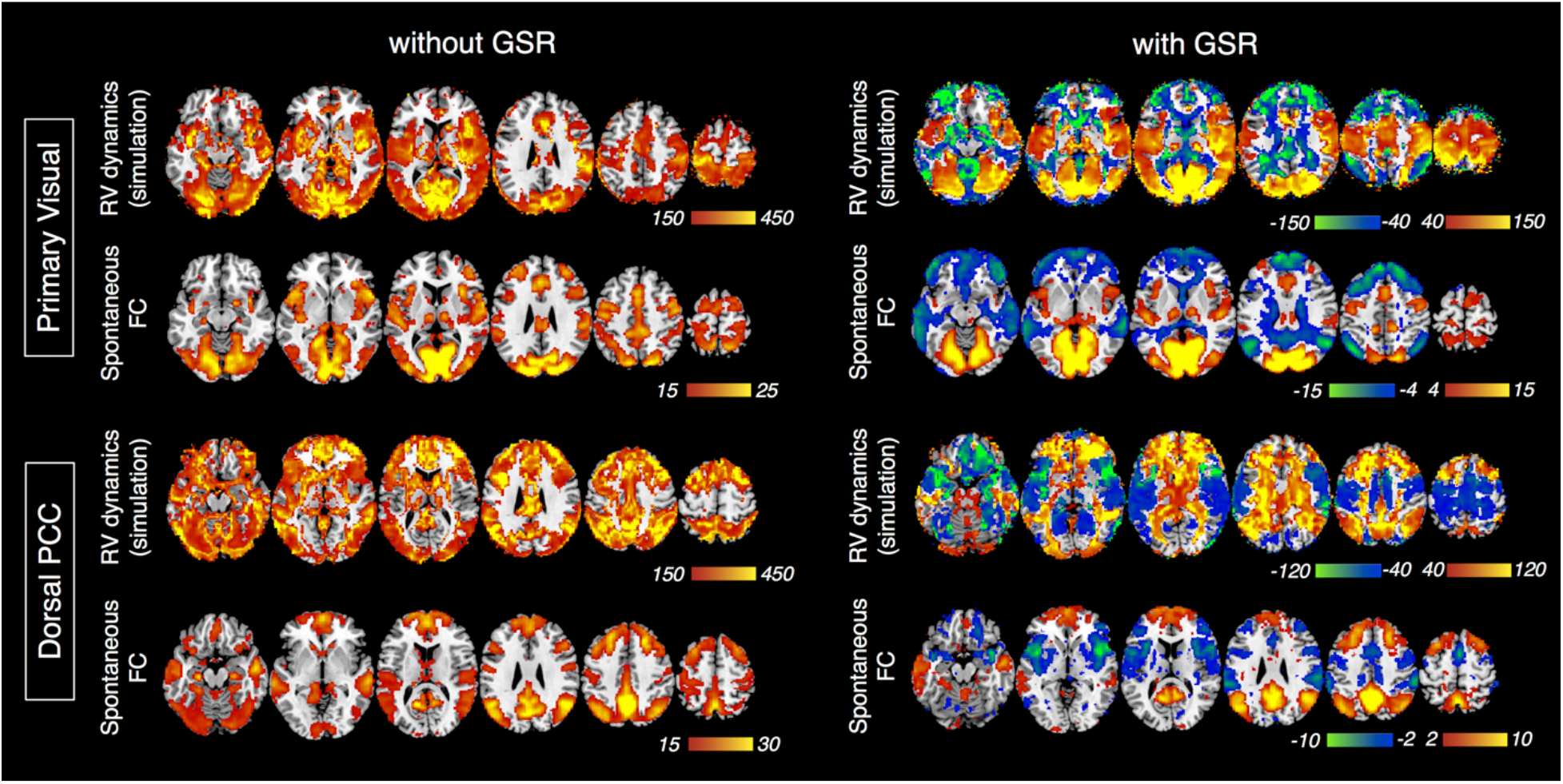
Group-level T-score maps of seed-based connectivity from “purely” RV-related dynamics and “clean” spontaneous fluctuations (derived from 190 synthesized and real datasets). The left column shows the results prior to GSR, and the right column shows the results after GSR. Note that the thresholds on the group-level T-scores of simulated RV dynamics were chosen to highlight most relevant regions rather than to make statistical inferences. No voxel clusters exhibited salient anti-correlations with either seed if GSR is not applied.

**Figure 5:**
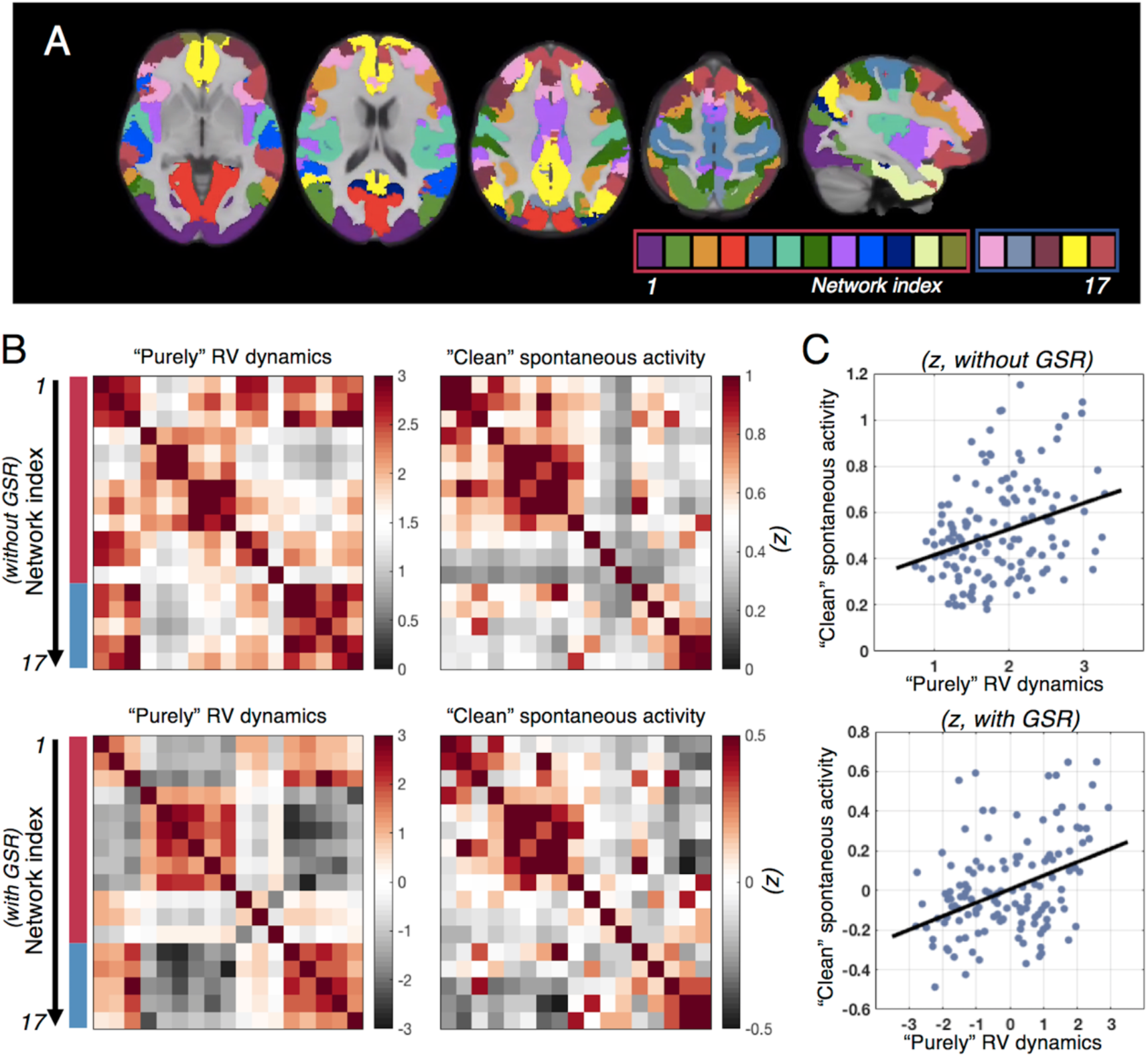
Results of network-based connectivity analysis applied to “purely” RV-related dynamics synthetic dataset and “clean” spontaneous fluctuations real dataset. (A) The Yeo et al. 17 RSN atlas (Yeo BT et al. 2011). Here we classify these RSNs into two groups, depending on whether they overlap more with the primary sensory RRF network (‘red box’) or with the frontoparietal RRF network (‘blue box’) identified in our data as shown in Fig. 3 (*k* = 2). (B) Z-scores (after Fisher transformation) of matrix connectivity of both “purely” RV dynamics and “clean” spontaneous dynamics, averaged across all subjects. Note that the colormap of the Z-score values differs between the two matrices; the range of Z-score values was adapted to visualize the matrix pattern. (C) Linear regression of “clean” spontaneous functional connectivity against “purely” RV connectivity (the averaged Z-scores shown in B, each dot represents one connectivity pair, 120 pairs in total).

As hypothesized, the GSR method, often used to remove systemic physiological noise contamination in resting-state fMRI data, could not remove the spatially-heterogenous physiological fluctuations observed in our data. Instead, this regression re-centered the correlation distributions and induced negative correlations between voxels in primary sensory cortices and in frontoparietal cortices (see Fig. 4 “purely” RV dynamics, with GSR, and Fig. 5B “purely” RV dynamics, with GSR, red vs. blue network groups’). Note that the unusually high T- and Z-scores seen in the apparent RV connectivity values (both with and without GSR) were caused by our assumption of subject-invariant RRFs and because we did not include any additional noise sources in the synthetic data (see Section 2.4.1).

Although we attempted to remove the physiological fluctuations from the resting-state fMRI data using model-based linear regression to obtain estimates of “clean” BOLD fluctuations reflecting spontaneous neuronal activity (see Section 2.4.2), clear similarities can be seen between the correlations derived from the synthetic dataset consisting of “purely” RV-related dynamics and measured dataset consisting of “clean” spontaneous fluctuations, both with and without GSR (Fig. 4, 5B). The similarity between these two datasets is further confirmed by the linear regression analysis of “clean” spontaneous activity against “purely” RV dynamics (120 pairs of network signals in Fig. 5B), which yielded *p* values of 4×1e^−4^ (*R^2^* = 0.11) without GSR and 1.1×1e^−6^ (*R^2^* = 0.16) with GSR (Fig. 5C). This apparent association between the “purely” RV-related dynamics and the “clean” spontaneous activity may be partly unexpected, however it is likely, in part, due to an incomplete removal of the physiological signals in the “cleaned” data and potentially also in part due to the underlying vascular anatomy and physiology shared by both of these datasets that influences both sets of BOLD fluctuations, as discussed in Section 4.2 below.

## 4 Discussion

### 4.1 General findings

Here, we show that BOLD dynamics responding to slow changes in respiratory volumes and heart rates exhibit heterogenous patterns across different brain regions. In the case of RRF dynamics, a clear distinction can be observed between primary sensory cortices and frontoparietal cortices; whereas in iCRF dynamics, the responses within the cerebral cortex are somewhat more homogeneous, although strong differences can be seen between the cerebral cortex and the rest of the brain including the cerebral white matter. Yet for both physiological responses, regional variability is evident, and these physiologically-driven dynamics can give rise to apparent “network” structures closely resembling common RSNs, regardless of whether systemic physiological denoising via GSR has been applied. It is worth highlighting that these apparent “physiological networks” stem primarily from spatially heterogenous response functions representing a BOLD signal change driven by some “physiological” effect, therefore they are distinct from conventionally-defined functional networks comprised of sets of segregated brain regions exhibiting synchronized fMRI dynamics presumably originating from coupled neuronal activity.

### 4.2 Potential explanations for the structured patterns of physiological responses

#### 4.2.1 Propagation pathways of RRF and iCRF dynamics

We observed that the fastest and also the largest RRFs were found in the brainstem, anterior cingulate cortex, thalamus and the primary sensory regions. Because of the limited temporal resolution of our fMRI data, we cannot discern the precise relative timing of the RRF between these regions. However, these observations agree with the well-characterized role of brainstem nuclei in regulating naturalistic breathing rhythms (Guyenet PG and DA Bayliss 2015; Ikeda K et al. 2017), as well as the well-known involvement of anterior cingulate cortex and thalamus in autonomic processes (Critchley HD et al. 2003; Beissner F et al. 2013).

The segregated patterns between the cerebral cortex and the rest of the brain (including the large vessels, white matter and ventricles) seen in the cardiac responses are akin to the coupling patterns between resting-state BOLD signals and peripheral measures of vascular tone reported previously (van Houdt PJ et al. 2010; Tong Y et al. 2013; Tong Y *et al*. 2017; Ozbay PS et al. 2018). In particular, the cardiac-related spatiotemporal dynamics observed here agree with the previously described propagation pattern of a low-frequency hemodynamic oscillation (measured via functional near-infrared spectroscopy and tracked through the brain using BOLD fMRI) reported by Tong and colleagues (Tong Y and BD Frederick 2010; Tong Y et al. 2013; Tong Y et al. 2017), in which these hemodynamic oscillations began in the largest feeding arteries then progressed up into the brain parenchyma and eventually appeared in the draining veins. This observed difference in the arrival time of the hemodynamic oscillations between the large arteries, cortex and large veins was conjectured to reflect the expected passage of blood “piped” through the brain (Tong Y *et al*. 2017). Although it is not trivial to interpret the direct relationship between the cardiac dynamics observed here and the hemodynamic oscillations reported in this prior work, some consistency in the spatiotemporal patterns can be expected given that they both presumably reflect changes in cardiac output regulated by the autonomic nervous system.

To evaluate whether the characterized physiological dynamics were also constrained by the vascular anatomy, we compared the spatial patterns of the strongest physiological responses with a recently published cerebrovascular atlas (Bernier M et al. 2018); we note that regions demonstrating strongest RRF/iCRF amplitudes indeed overlap with regions with greater likelihood of high vessel density (Fig. 6, ‘Regions with max. RRF/iCRF amp.’ with ‘Arterial/Venous vessel density’, highlighted by purple rectangles). However, we were also able to identify multiple regions with high vascular density but low BOLD amplitudes relative to other regions (highlighted by green rectangles), suggesting that vascular anatomy may not be the only factor that determines the observed physiological dynamics.

**Figure 6:**
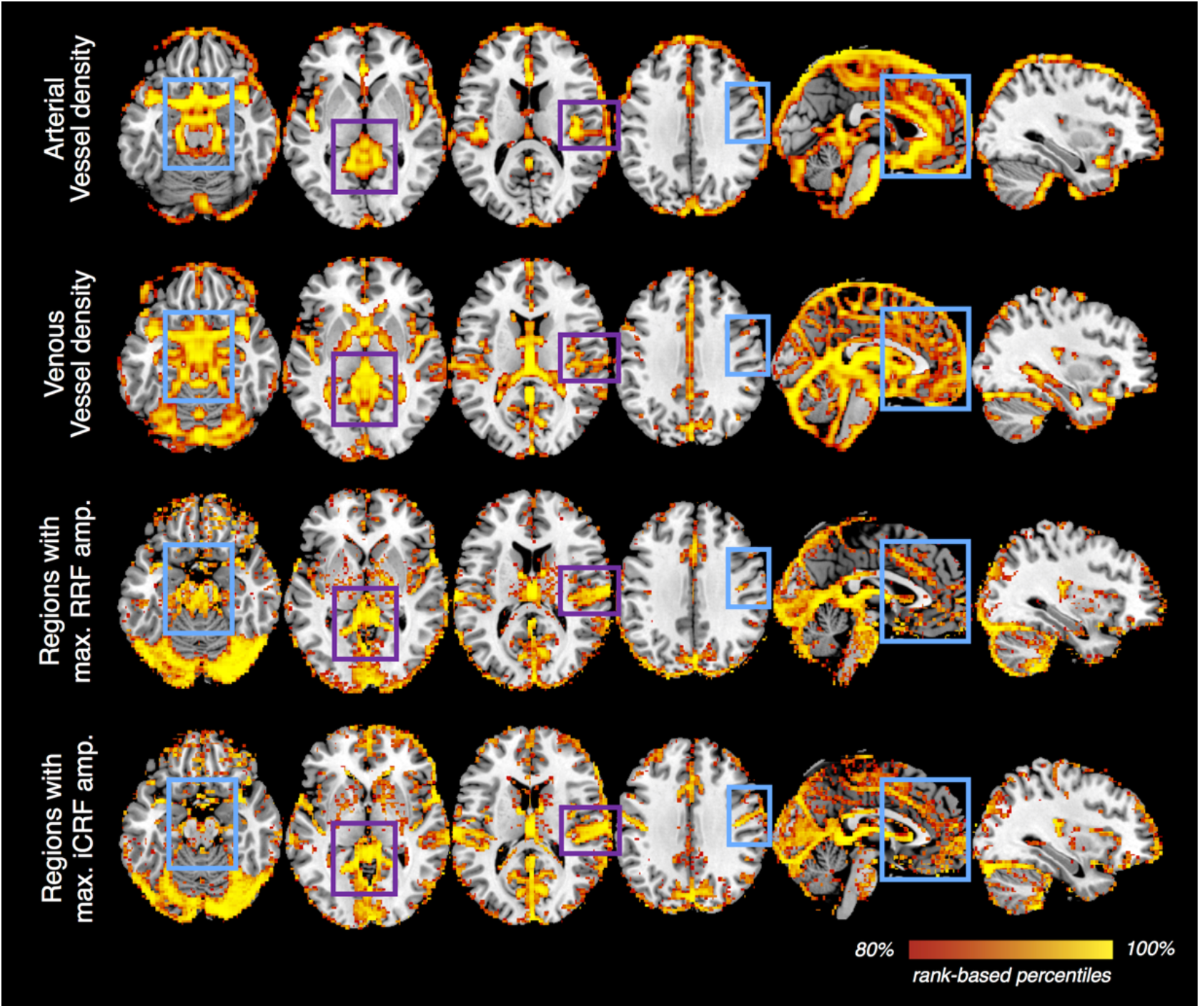
Vessel density vs. the amplitudes of voxel-wise RRFs/iCRFs. A recently published atlas of cerebral vessels (the probability map of local vesselness, (Bernier M et al. 2018)) was moderately smoothed (with an isotropic 3D Gaussian kernel, FWHM = 5 mm) to serve as an approximate estimate of local vessel density for both the arterial and venous compartments of the intracranial vasculature. For each brain voxel, the maximum value of RRF waveform and the minimal value of iCRF waveform were calculated to reflect the amplitudes of RRFs/iCRFs at each brain location (without spatial smoothing). For each map shown, voxels within the top 20% percentiles (based on the probability values provided in the arterial/venous vessel density map and amplitudes in RRF/iCRF) are displayed to provide an appropriate comparison across the different quantities. This qualitative comparison shows that vascular density may indeed explain some regional variability in the observed slow physiological dynamics (e.g., regions highlighted by purple rectangles), however, it is unlikely to be the only factor that determines the spatiotemporal patterns of RRFs or iCRFs (e.g., regions highlighted in cyan rectangles).

#### 4.2.2 Shared spatial patterns between “physiological” dynamics and spontaneous functional activity

To interpret the observed similarity between the putative “physiological networks” and conventional RSNs, we must consider possible limitations of our methodology. Previous studies have cautioned that the subspace spanned by a large number of regressors (for instance, the 10 regressors used here) inevitably will remove variance of meaningful, neuronally-relevant functional information contained within the raw BOLD time-series data, even if the regressors were derived from physiological recordings based on external sensors (Bright MG and K Murphy 2015; Chen JE et al. 2017). We addressed this concern with a control analysis based on repeating the regression with an identical number of regressors with RV/HBI measures from a different subject, which only produced responses with negligible intensity fluctuations, suggesting that this potential caveat may not apply to our findings. Furthermore, we took the additional step of synthesizing new datasets containing only RV dynamics to test whether the observed network patterns could be formed purely by slow respiratory fluctuations (described in Section 2.4), rather than considering only the variance fitted by physiological bases (described in Section 2.2), which also circumvented this potential caveat.

A second possibility we considered relates to the common vascular origins that underlie both “physiological” and neural signals. RRFs/iCRFs may share similar temporal properties with neuronally-driven hemodynamic response functions (HRFs), because both waveforms reflect hemodynamic changes directly following altered blood flow and oxygen consumption through local vasculature. Qualitatively, some relationship can be seen between local vascular anatomy and density and the “physiological” responses (see Fig. 6), and vascular anatomy and density also appears to have some influence on spontaneous neuronal activity (Tak S et al. 2014; Vigneau-Roy N et al. 2014; Tak S et al. 2015). Direct evidence supporting this argument has been offered by studies showing that the vascular response latency identified by a breath-hold task closely resembled the task-evoked hemodynamic delay patterns in primary sensory regions (Chang C et al. 2008; Li Y et al. 2018). Along similar lines, a recent study observed a marked reduction in the strength of resting-state connectivity if regional HRF shapes were taken into account through blind de-convolution (Rangaprakash D et al. 2018), which indeed implies a possible link between regionally varying HRFs and apparent RSN structures reported in the literature, similar to how RRFs/iCRFs determine apparent “physiological” networks reported here.

Another possible, and more speculative, interpretation may be found by presuming a tight coupling between vascular regulation and neuronal activity, as suggested in a recent study^1^. These “physiological networks” characterized in the present study, and in the study of Bright et al. using a hypercapnia challenge, all point to mechanisms of vascular regulation in the brain that can promptly provide fast and efficient blood delivery to multiple, remote brain regions that tend to co-activate due to similarities in their neuronal function or to direct neuronal coupling, resulting in coherences in vascular fluctuations that in some way reflect similarities in neuronal function. Such large-scale vascular regulatory architecture may develop concurrently with large-scale neuronal pathways established during development (Quaegebeur A et al. 2011), and reshaped/remodeled during learning and experience (Black JE et al. 1990). In this sense, the “physiological networks” may reflect neuronal networks because the vasculature is adapted to the neuronal network structure.

### 4.3 Implications for functional connectivity studies

Having recognized that apparent “physiological connectivity” shares overlapped information with spontaneous functional activities, it follows naturally to query the extent to which such “physiological connectivity” influences the overall connectivity estimates during resting state. Several studies have demonstrated that RV and heart rate fluctuations can account for considerable proportions of signal variance in standard-resolution fMRI data which are prone to be physiological-noise dominated ((Birn RM et al. 2006; Chang C et al. 2009; Golestani AM et al. 2015), and illustrated in Supplementary Fig. S3). We noticed that even in the modern-resolution HCP fMRI data considered here, which, due to their smaller voxel sizes (Triantafyllou C et al. 2005; Triantafyllou C et al. 2011), will exhibit less physiological noise dominance than standard-resolution fMRI data, a strikingly strong coupling was observed between the BOLD time-series data and the RV fluctuations in multiple subjects. In these cases, “physiological connectivity” may dominate the overall apparent connectivity estimates. Traces of resting-state BOLD data from an exemplar subject are illustrated in Fig. 7A (signals in most gray matter regions tracked RV, and the coupling was most evident during the period highlighted in a gray box). Notably, despite the similarity of the resting-state BOLD fluctuations with the RV changes seen globally across the brain, BOLD signals measured within each ROI taken from the Yeo 17-network parcellation exhibit time delays and other differences in temporal properties relative to one another (Fig. 7A, C, ‘Yeo 17-network signals’, comparing different colors). As has been cautioned by our earlier analyses, such regional differences in the temporal properties of the BOLD signal—albeit with a consistent coupling with the slow respiratory fluctuation—will likely drive a decomposition of these regional signals in an ICA. Indeed, an ICA performed on the whole-brain BOLD data during this period of slow respiratory fluctuations yielded multiple RSN-like, but strongly RV-coupled spatial networks (supplementary Fig. S4), and the corresponding ICs exhibited similar ranges of time delays and other differences in temporal properties relative to one another (Fig. 7A, C, ‘IC signals’; the subset of data exhibiting the strongest coupling with the RV waveform (highlighted in the gray box) were included for ICA). This example offers an illustration that the influence of “physiological connectivity” can be very substantial.

**Figure 7:**
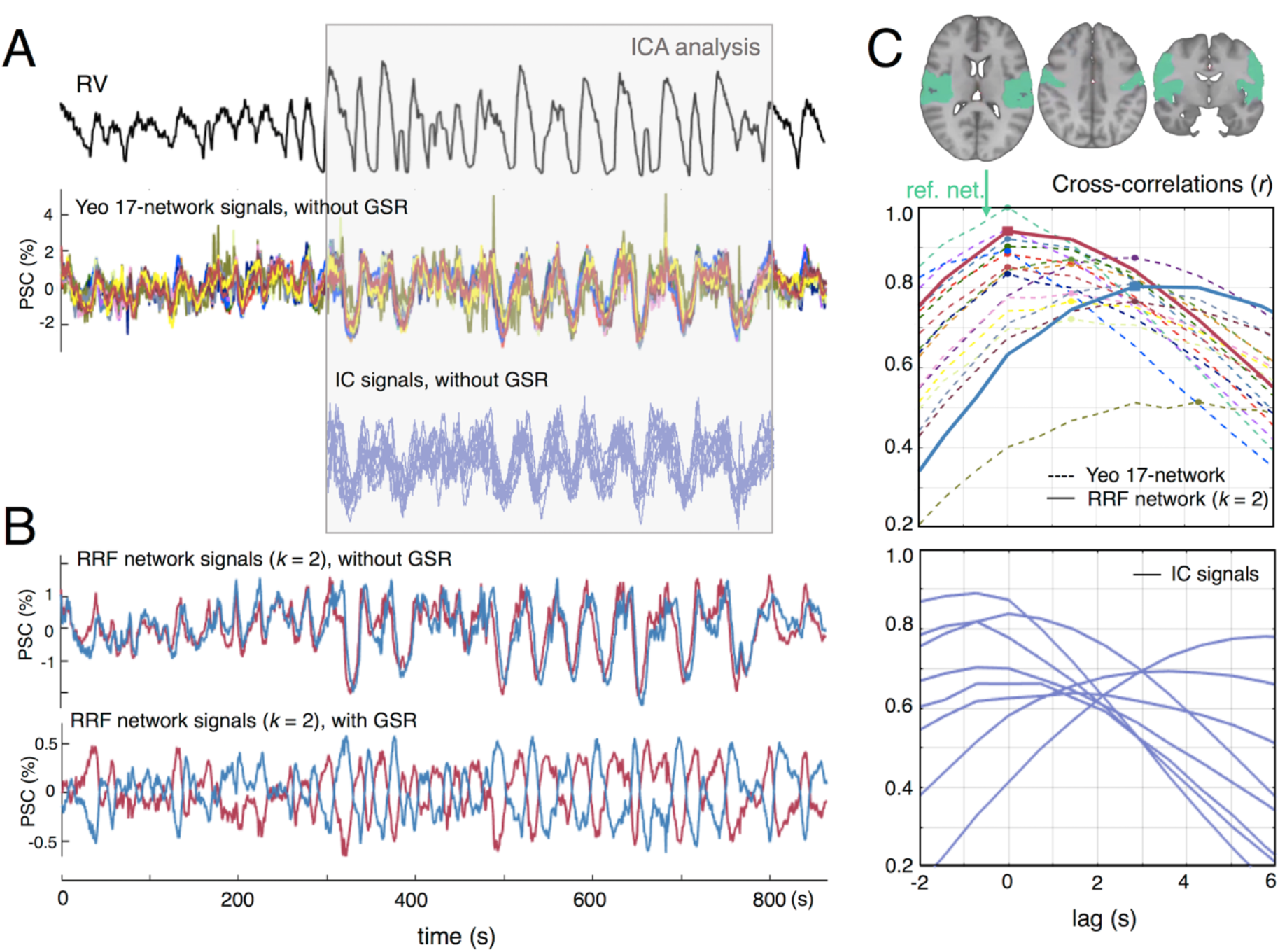
Data of an exemplar subject (ID: 102008) with substantial physiological effects in the total BOLD fluctuations are shown. (A) Tight coupling between the RV waveform and widespread BOLD signals was observed (‘Yeo 17-network signals’, PSC: percent signal change). Due to the spatial heterogeneity of BOLD fluctuations correlated with RV, ICA resolved multiple components with time courses closely resembling lagged versions of the RV waveforms (‘IC signals, without GSR’, each line represents one independent component (IC), and the corresponding spatial maps of different ICs are shown in Supplementary Fig. S4, which resemble several common RSNs). (B) In agreement with our RRF results (Fig. 3 ‘RRF networks, *k*=2’), signals averaged within the primary sensory cluster (red trace) preceded those averaged within the frontoparietal cluster (blue trace). GSR only removed common variance present in the data, leaving an anti-correlated pair of residual signals. (C) Summary of temporal lags computed among all the RV-like BOLD dynamics. This was obtained by computing the cross-correlations between all fMRI signals within the Yeo 17-network (top), and from the results of the ICA performed on the time interval indicated in panel A (bottom). For the lags between the Yeo 17-network components, the solid lines represent the primary sensory cluster (red trace) and the frontoparietal cluster (blue trace), and the dashed lines represent the remaining 15 networks with line colors matching the convention used in Fig. 5. All lags are calculated relative to a reference network signal (‘ref. net.’), a network from the Yeo atlas that comprised of auditory and sensorimotor regions, and the ROI of this reference network (in green colors) is displayed above the plots.

Given the significant contribution from “physiological” effects, the question arises of how we can separate spontaneous neuronal fluctuations of interest from these slow “physiological” dynamics to remove these noise sources. If we simply treat these “physiological” fluctuations as a nuisance, as is common across the majority of resting-state studies, it would be preferable to include voxel-specific regressors that can address spatially heterogeneous RRF/iCRF patterns (e.g., by modeling physiological regressors with possible temporal or dispersive derivatives) rather than regressing out a single waveform, e.g., using the response modelled by the canonical response functions without including temporal or dispersive derivatives, or the global signal. It is beyond the scope of this study to argue the advantages or disadvantages of GSR, which have sparked many discussions but have not yet reached at a consensus (Liu TT et al. 2017; Murphy K and MD Fox 2017). Instead, we simply want to caution the possibility that, in addition to the well-known caveat regarding the appearance of anti-correlations following GSR (i.e., mandating all correlations summing to a non-positive value (Murphy K et al. 2009)), GSR and other single-waveform regression approaches may not minimize region-wise “physiological” fluctuations sufficiently, as has been previously argued (Power JD et al. 2018). Differences between local “physiological” dynamics (temporal lags or slight shape alterations) could be amplified by GSR due to the removal of common variance (illustrated in Fig. 7B), yet still generate network-like structures that could be mistaken for, or obscure, actual coupled neuronal fluctuations (evidenced by the group-level results post GSR shown in Figs. 4 and 5).

Surprisingly, beyond the context of resting-state functional connectivity, possible physiological contributions have received much less attention in studies examining altered brain states. Given that we have demonstrated apparent “connecting” clusters may arise even in the absence of actual neuronal segregations among different cortical areas, our findings advise caution in interpreting connectivity results in non-resting brain conditions that are associated with notable changes in brain physiology and possibly higher fractional contribution from these “physiological” dynamics. This includes, for instance, different stages of sleep (Webb P 1974; Hudgel DW et al. 1984; Buchsbaum MS et al. 1989; Boyle PJ et al. 1994; Bonnet MH and DL Arand 1997; Nofzinger EA et al. 2002) and anesthesia (Dripps RD and JW Severinghaus 1955; Fink BR et al. 1962; Latson TW et al. 1992; Alkire MT et al. 1995; Alkire MT et al. 1997).

Beyond the discussion of physiological dynamics, apparent connectivity may theoretically be caused by spatially heterogeneous responses (lags and shapes) elicited by any systemic changes when applying traditional FC analyses (such as seed-based correlation, ICA or spatial clustering), which implicitly assume the network or brain-state of interest is fixed or static either over time or space. Apart from the physiological dynamics characterized here, recent studies have also identified—with purely data-driven approaches—several temporally and spatially evolving patterns that co-exist during the resting state, and have shown that these evolving spatiotemporal patterns account for considerable variance of the overall BOLD fluctuations (Majeed W et al. 2011; Mitra A et al. 2015; Byrge L and DP Kennedy 2018; Abbas A et al. 2019). Identifying the causes of these evolving patterns (e.g., physiology, neural activity or beyond) may point to a valuable avenue for discerning the nature of spontaneous brain FC during resting state.

### 4.4 Further technical considerations and future directions

In the current analysis, voxel-wise physiological responses were fitted with a set of pre-defined basis functions (Fig. 1C), therefore yielding a less flexible method to estimate response waveforms compared to the finite impulse response (FIR) based de-convolution (i.e., using a set of temporally delayed delta functions to fit the response). We did not adopt the latter approach for this study because this is expected to generate extremely noisy estimates in the presence of strong temporal autocorrelations in both RV and HBI waveforms (as both were derived across 6-s-long sliding windows centered at each fMRI time sample point) (Birn RM *et al*. 2008). A potential future improvement could be achieved by performing an FIR-based approach, but incorporating moderate regularization, e.g., smoothness (Chang C *et al*. 2009), to suppress noise enhancement from the ill-conditioned deconvolution.

In agreement with a recent study^2^ that applied a novel algorithm to de-convolve the physiological recordings from the resting-state BOLD data to estimate global respiratory and cardiac response functions in 41 HCP subjects, we also observed a faster global RRF and CRF (the inverse of iCRF) than the response functions reported previously by Birn et al. (2008) and by Chang et al. (2009). Multiple possibilities may explain the observed differences with these earlier studies. Firstly, in lieu of averaging BOLD responses elicited by a series of cued deep breaths (which are strong, evoked changes in respiration depth and rate) as performed by Birn et al., we used resting-state RV (a temporally smooth estimate of natural, instantaneous respiratory dynamics) during free breathing to characterize RRFs, and may therefore observe a slightly faster RRF. Secondly, RV and heart rate (inverse of HBI) reported by Chang et al. were sampled at longer temporal sampling rates (equivalent to the TR value of the fMRI acquisition, 2 s, which is about three times the sampling interval of the fMRI data used in the present study), and correspondingly, the resulting RRF or CRF should equal the temporal averaging of our RRFs/CRFs across three consecutive TRs. Thirdly, the RRF and CRF reported by Chang et al. were derived using FIR-based de-convolution with Gaussian temporal smoothness priors, which yielded smoother response functions but may also filter out potentially faster responses. Nonetheless, such differences suggest that although RRFs/iCRFs characterized here can portray the dynamics of different physiological processes, the exact timing may vary slightly with the fMRI acquisition protocols and data analyses strategies.

While we have focused on the common physiological pattern observed across a large cohort of subjects, we also noticed considerable inter-individual variability in the RRFs and iCRFs (shown in Supplementary Fig. S5), in support of previous proposals advocating for subject-specific, global RRFs and CRFs (Falahpour M et al. 2013; Cordes D et al. 2014; de la Cruz F et al. 2017) to single-voxel levels. A possible future direction will be to identify the role of subject-specific physiological dynamics in the across-subject variability of apparent FC estimates, which may be important for both interpreting single-subject level connectome fingerprinting (Finn ES et al. 2015), and for improving the group-level sensitivity of fMRI as either a neural or physiological biomarker in future studies.

Recent studies have also suggested that respiratory changes are coupled with noticeable displacements in head motion measurements (Power JD et al. 2017; Byrge L and DP Kennedy 2018; Power JD *et al*. 2018), and it is thus possible that the resolved respiratory patterns in the fMRI data may partially reflect apparent head movements. However, motion is not likely to be the primary explanation for our findings, because the detected spatial patterns in the physiological responses closely trace anatomical structures within the brain (Fig. 2A), and that these “physiological” networks (Fig. 3) differ from those spurious local or long-range correlation patterns characterized in previous studies investigating the impact of head motion of functional connectivity estimates (Power JD et al. 2012; Satterthwaite TD et al. 2017). Furthermore, it is non-trivial to interpret motion estimates at instances of dramatic respiratory changes, as they may arise from magnetic field perturbations induced by respiration, instead of mechanical displacements of the head (Van de Moortele PF et al. 2002; Fair DA et al. 2018; Chen JE et al. 2019). Nevertheless, to address the concern of possible motion-driven fluctuations in the observed physiological dynamics, we separated the subjects into two cohorts based on the extent of correlation between motion estimates and RV or HBI measures. No notable differences in the spatiotemporal patterns of RRFs and iCRFs can be identified between these two groups except for response intensities (see Supplementary Figs. S6 and S7), suggesting that it is unlikely that head motion contributes to the patterns of physiological responses seen in our resting-state BOLD fMRI data.

Although the measured response functions were time-locked to RV or HBI changes, they may not necessarily represent BOLD changes directly caused by these two specific forms of physiological signals. First, other systemic changes, e.g., changes in blood pressure (Whittaker J et al. 2018) or other changes in the cardiac signal (such as pulse height (van Houdt PJ et al. 2010; Tong Y et al. 2013; Tong Y *et al*. 2017; Ozbay PS et al. 2018)), could also elicit notable BOLD fluctuations that are difficult to isolate from RV or HBI dynamics due to the expected interdependence among these different aspects of physiological dynamics. Second, changes in systemic physiology may reflect changes in brain state, which will also trigger either local or large-scale neuronally-driven hemodynamic responses that co-occur with the associated physiological responses (Yuan H et al. 2013). Particularly during relatively long resting-state scans (15 minutes in the HCP protocol), subjects may enter into low-vigilance states, or even gradually descend into light sleep (Glasser MF et al. 2018), in which a global increase in BOLD fluctuations would occur (Fukunaga M et al. 2006; Wong CW et al. 2013). The RRFs and iCRFs characterized in this study may therefore, in part, reflect BOLD changes related to changes in arousal level. It is challenging to disentangle the relative contributions from different physiological or neuronal mechanisms based on fMRI signatures alone; concurrent monitoring of more complete sets of physiological changes, including concurrent measures of arousal level, and acquisition of more direct neural (or metabolic) ground truth may aid differentiation among various mechanisms in future studies.

Finally, while most studies consider neuronal networks as the primary interest, such that these “physiological networks” are confounds as discussed above, there is also an emerging field of “physiological MRI” focused on using fMRI techniques to characterize vascular physiology in patients with various lesions or neurodegenerative diseases associated with vascular dysfunction (Golestani AM et al. 2016; Jahanian H et al. 2017; Lu H 2019). In such applications, “physiological networks” discussed here may provide meaningful characterizations of blood flow regulation and vascular anatomy, and hold great potential in serving as clinical biomarkers that are complementary to neuronal networks.

## 5 Conclusions

In this study, we have characterized regional respiratory and cardiac response functions based on modern fast-fMRI acquisitions. We show that the spatial heterogeneity of these physiological responses in the BOLD data can give rise to correlated signals across diverse brain regions that resemble well-known large-scale brain networks, and that the influence of physiological dynamics on spontaneous brain fluctuations cannot be eliminated using a single-waveform-based regression approach, e.g., GSR. These “physiological” networks characterized in this study may themselves provide meaningful characterizations of blood flow regulation and vascular anatomy and their influences on BOLD dynamics. However, given that apparent connectivity can arise from systemic physiological changes alone, our findings also advise caution in interpreting apparent connectivity in future studies that examine brain states or populations with altered or atypical brain physiology.

## Funding

This work was supported in part by the National Institute of Biomedical Imaging and Bioengineering at the National Institutes of Health (grant numbers P41-EB015896, R01-EB019437), by the *BRAIN Initiative* (National Institute of Mental Health at the National Institute of Health: grant numbers R01-MH111438, R01-MH111419), by the National Institute of Mental Health at the National Institute of Health (grant number R00-MH111748), by the National Institute of Environmental Health Sciences at the National Institute of Health (grant number K22ES028048), and by the MGH/HST Athinoula A. Martinos Center for Biomedical Imaging; and was made possible by the resources provided by NIH Shared Instrumentation grants S10-RR023043 and S10-RR019371. Data were provided by the Human Connectome Project, WU-UMinn Consortium (Principal Investigators: David Van Essen and Kamil Ugurbil; U54-MH091657) funded by the 16 Institutes and Centers at the National Institute of Health that support the NIH Blueprint for Neuroscience Research; and by the McDonnell Center for Systems Neuroscience at Washington University.

## Acknowledgements

The authors acknowledge Drs. Vitaly Napadow, Nicola Toschi, Roberta Sclocco and Andrea Duggento for valuable discussions regarding the present results.

## Supplementary Material

**Fig. S1:**
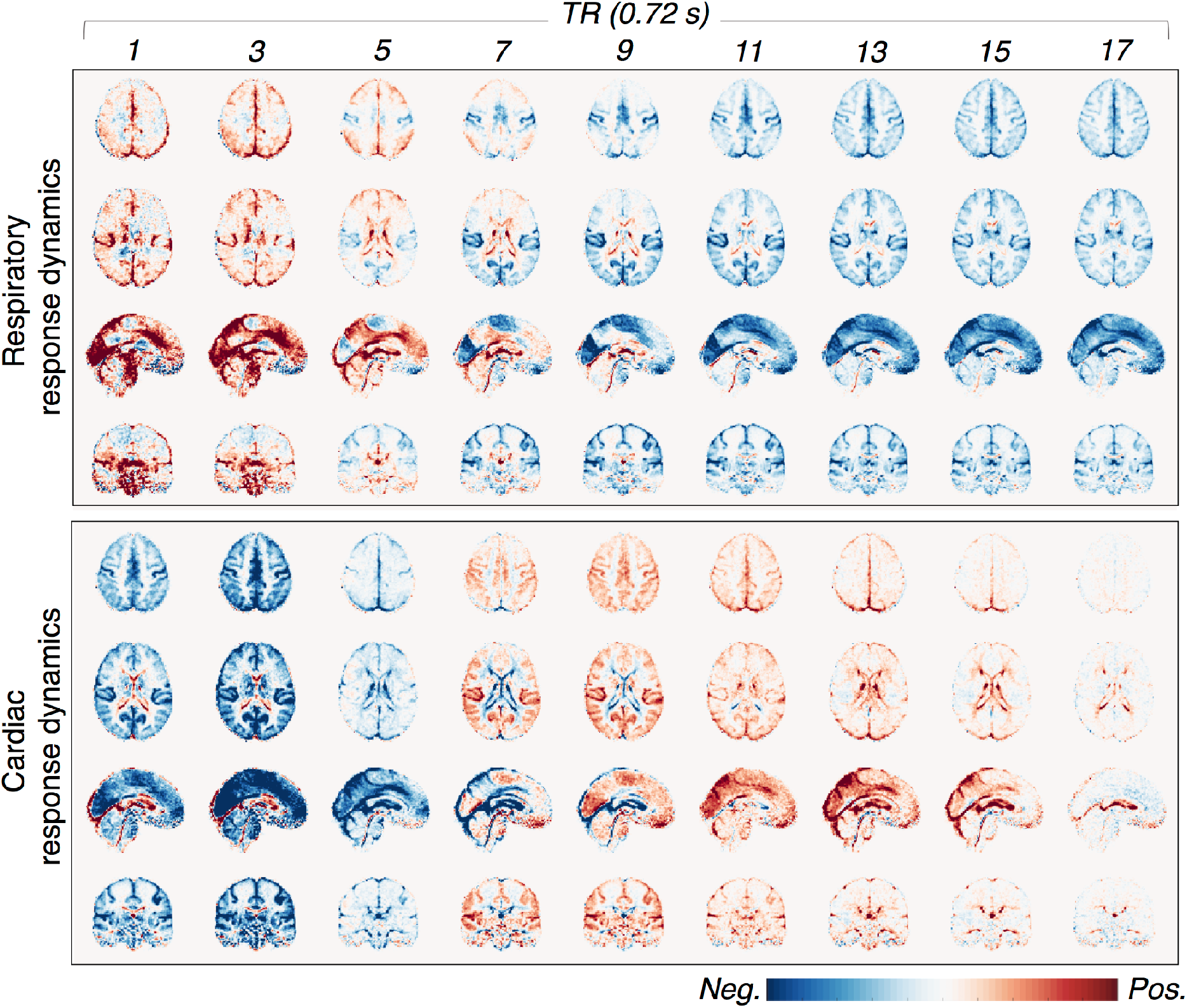
Spatiotemporal dynamics of voxel-wise RRFs and iCRFs averaged across 190 subjects, without spatial smoothing.

**Fig. S2:**
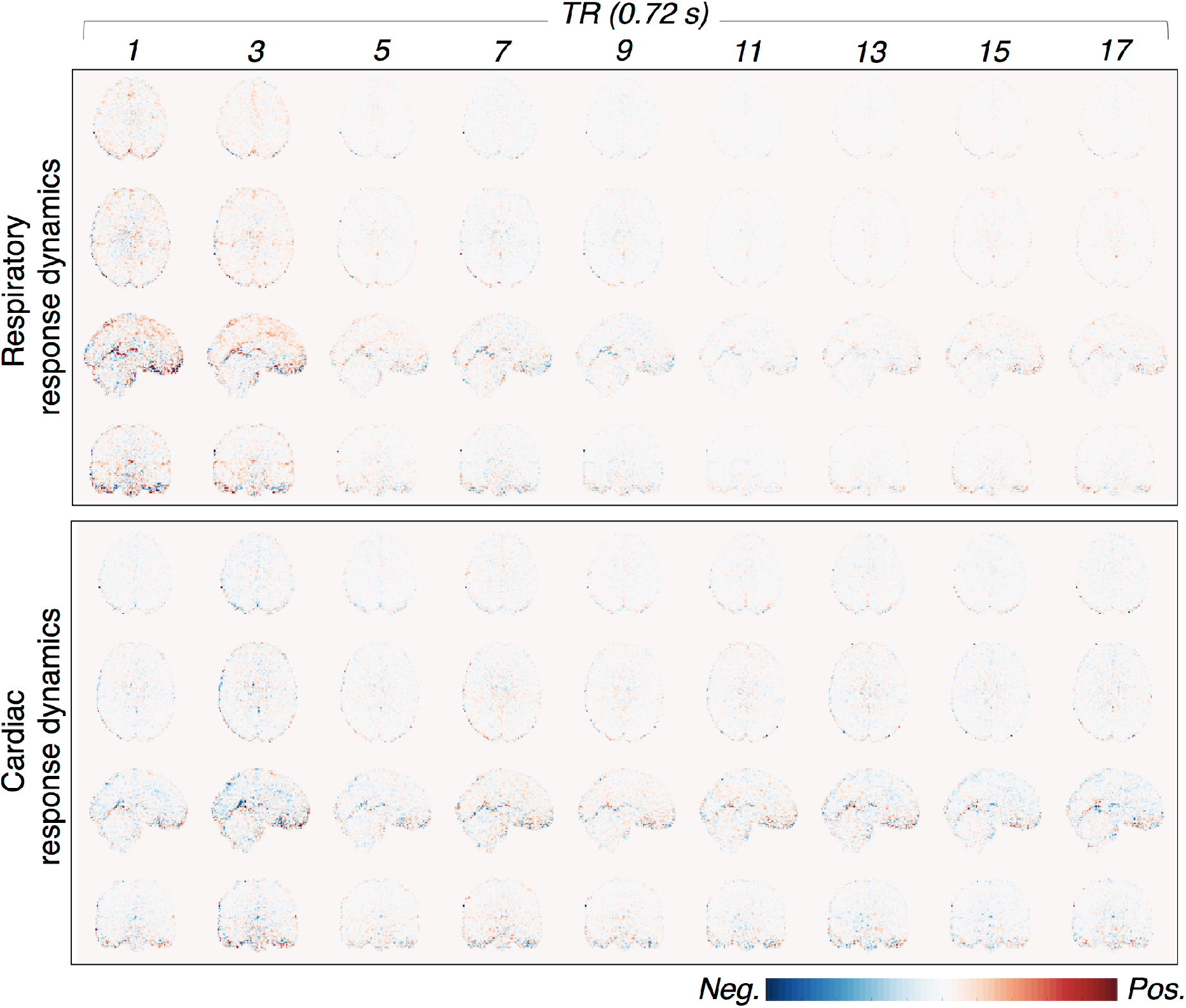
Results of control analysis consisting of deriving voxel-wise BOLD response function in each subject from RV/HBI waveforms from a different subject, and then averaged across all 190 subjects, without spatial smoothing. The displayed color range is identical with Fig. 2A and Fig. S1.

**Fig. S3:**
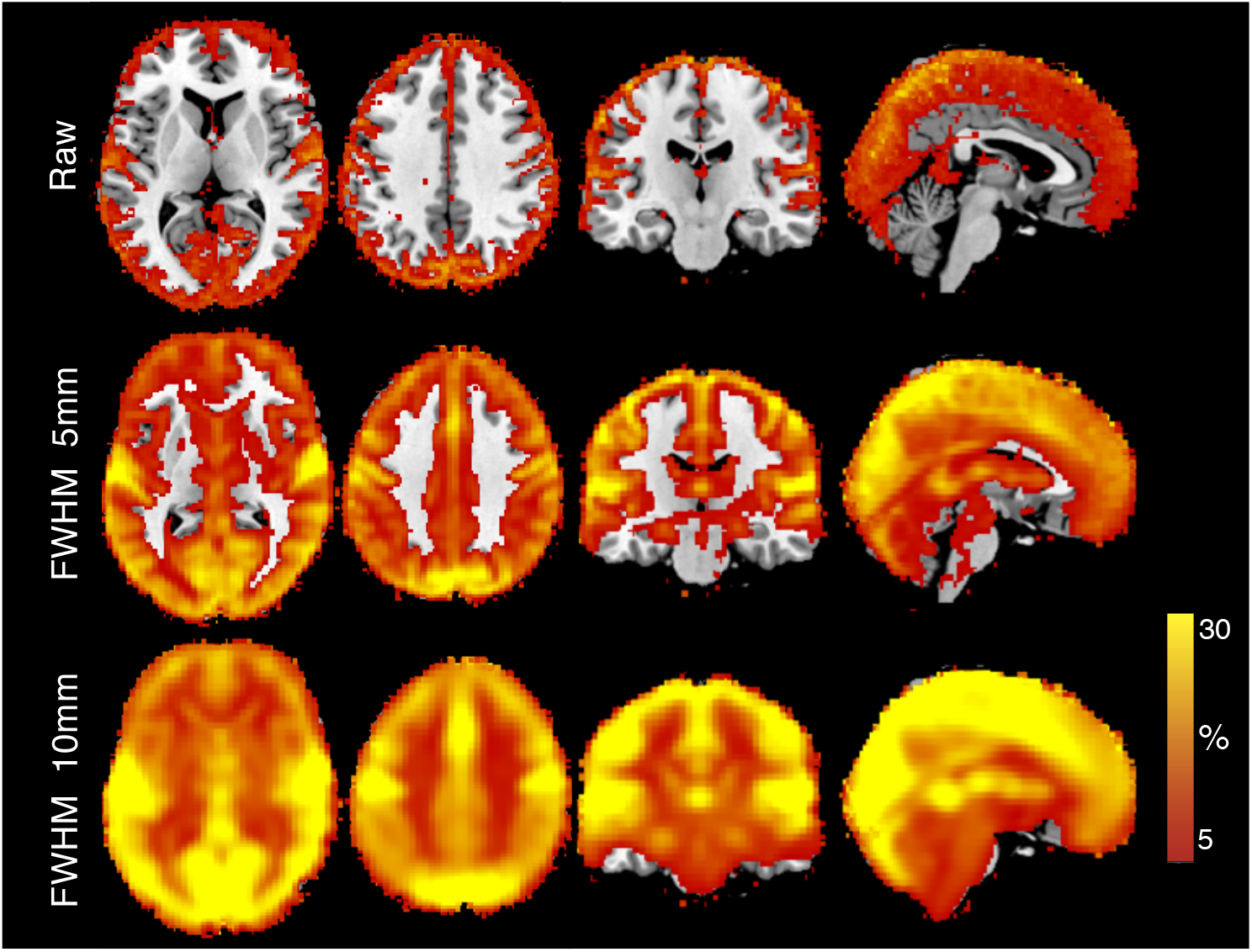
Fractional contribution from slow RV and HBI fluctuations as a function of spatial resolution (manipulated by explicit spatial smoothing with different kernel sizes). This was calculated by evaluating the voxel-wise variance explained by the modeled physiological fluctuations (i.e., the basis set of responses shown in Fig. 1C), then averaging across 190 subjects.

**Fig. S4:**
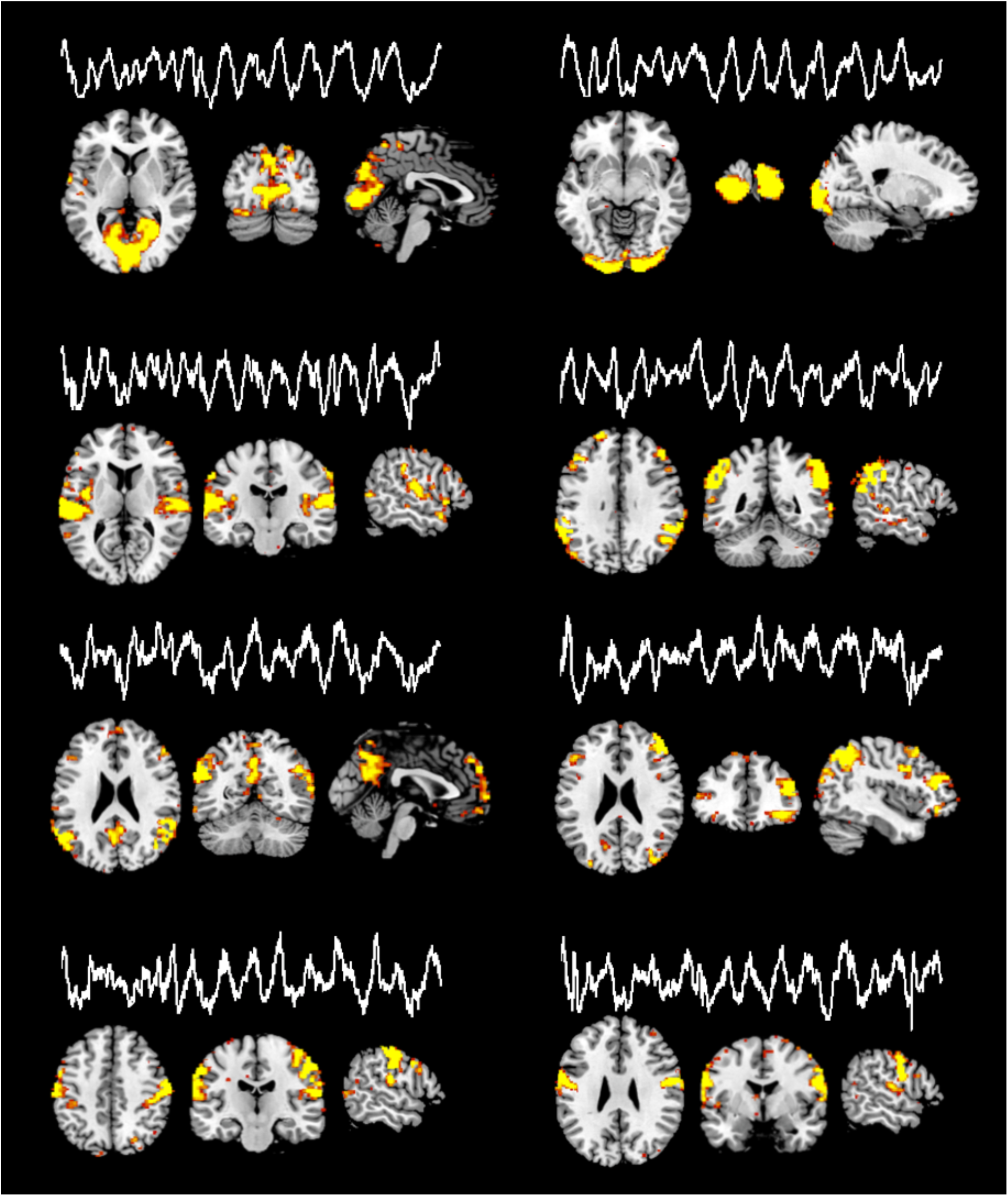
Spatial maps and time courses of the 8 RSN-like independent components (ICs) shown in Fig. 7. This was derived using the data of a single HCP subject (ID: 102008, with spatial smoothing, FWHM = 5 mm, time interval: 300–800 s). ICA was performed using FSL MELODIC, and the number of ICs was estimated automatically.

**Fig. S5:**
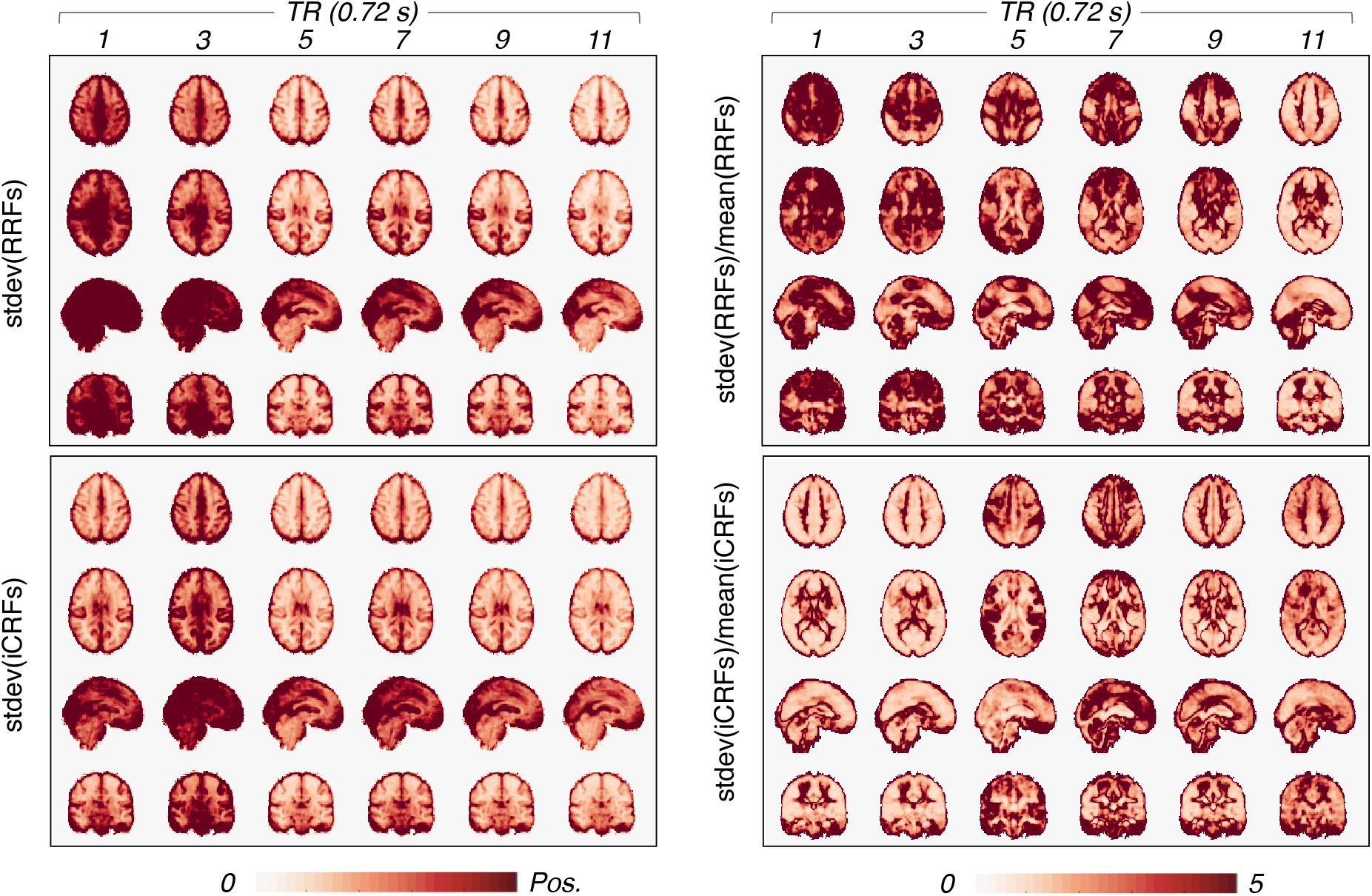
Considerable inter-subject variability in RRF and iCRF patterns. Left column: standard deviation of RRF and iCRF intensities across 190 subjects (the displayed color range is identical with Fig. 2A); right column: the standard deviations are normalized by the mean RRF and iCRF intensities (i.e., Fig. 2A).

**Fig. S6:**
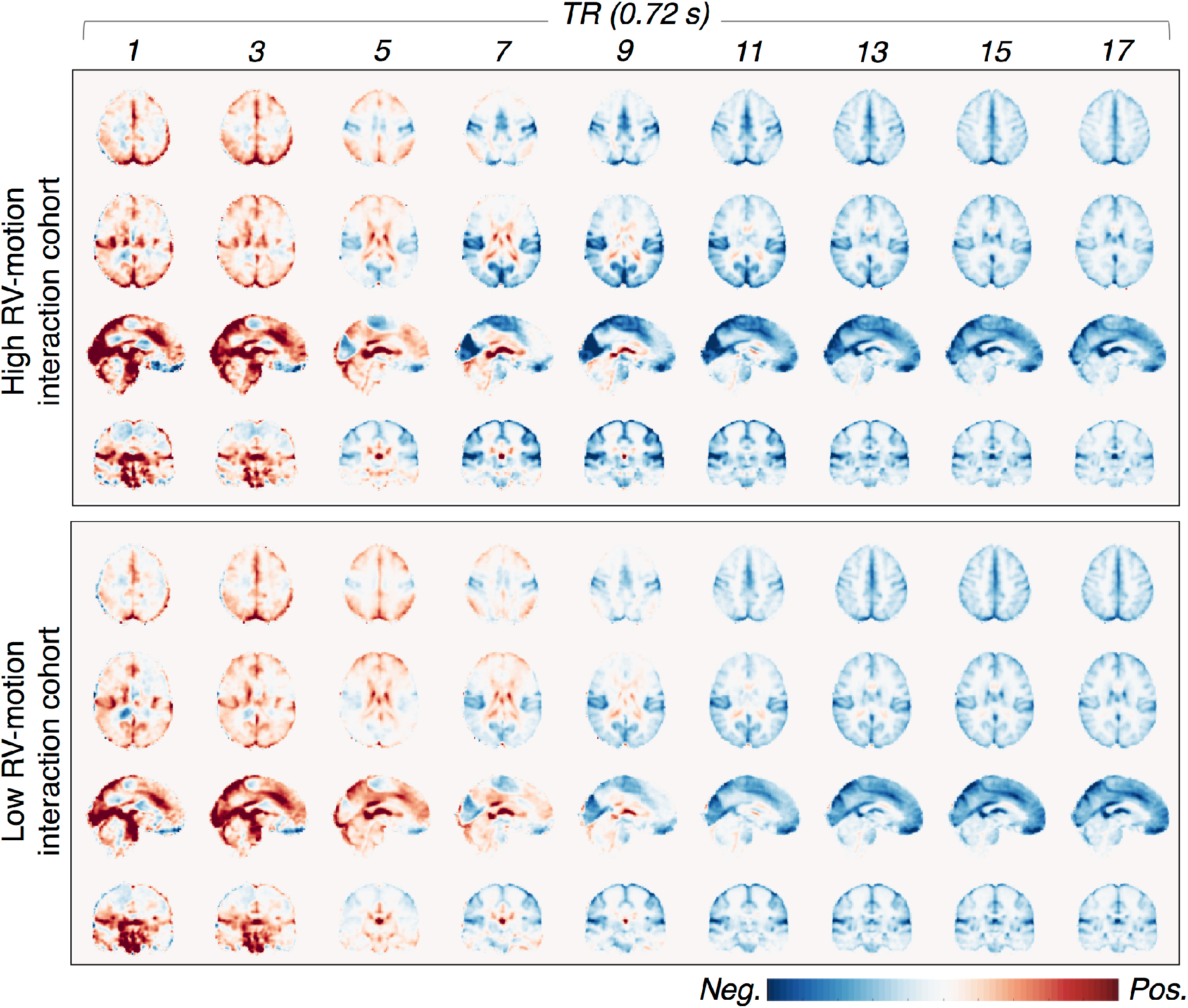
Influence of “head motion” on RRF and iCRF dynamics. Subjects were separated into two cohorts according to the interaction between head motion and RV measurements, quantified as the absolute value of the linear Pearson correlation between the RMS of head movements (‘*Movement_AbsoluteRMS.txt*’ released by HCP) and RV changes, |*r*_RV-motion_|. RRFs were averaged within two separate cohorts of subjects with different levels of RV-motion interactions: ‘high RV-motion interaction cohort’ includes the 95 subjects with higher |*r*_RV-motion_| values, mean±stdev = 0.24±0.10; and ‘low RV-motion interaction cohort’ includes the 95 subjects with lower |*r*_RV-motion_| values, mean±stdev is 0.054 0.039. No prominent differences can be observed between these two groups except for changes in response intensities.

**Fig. S7:**
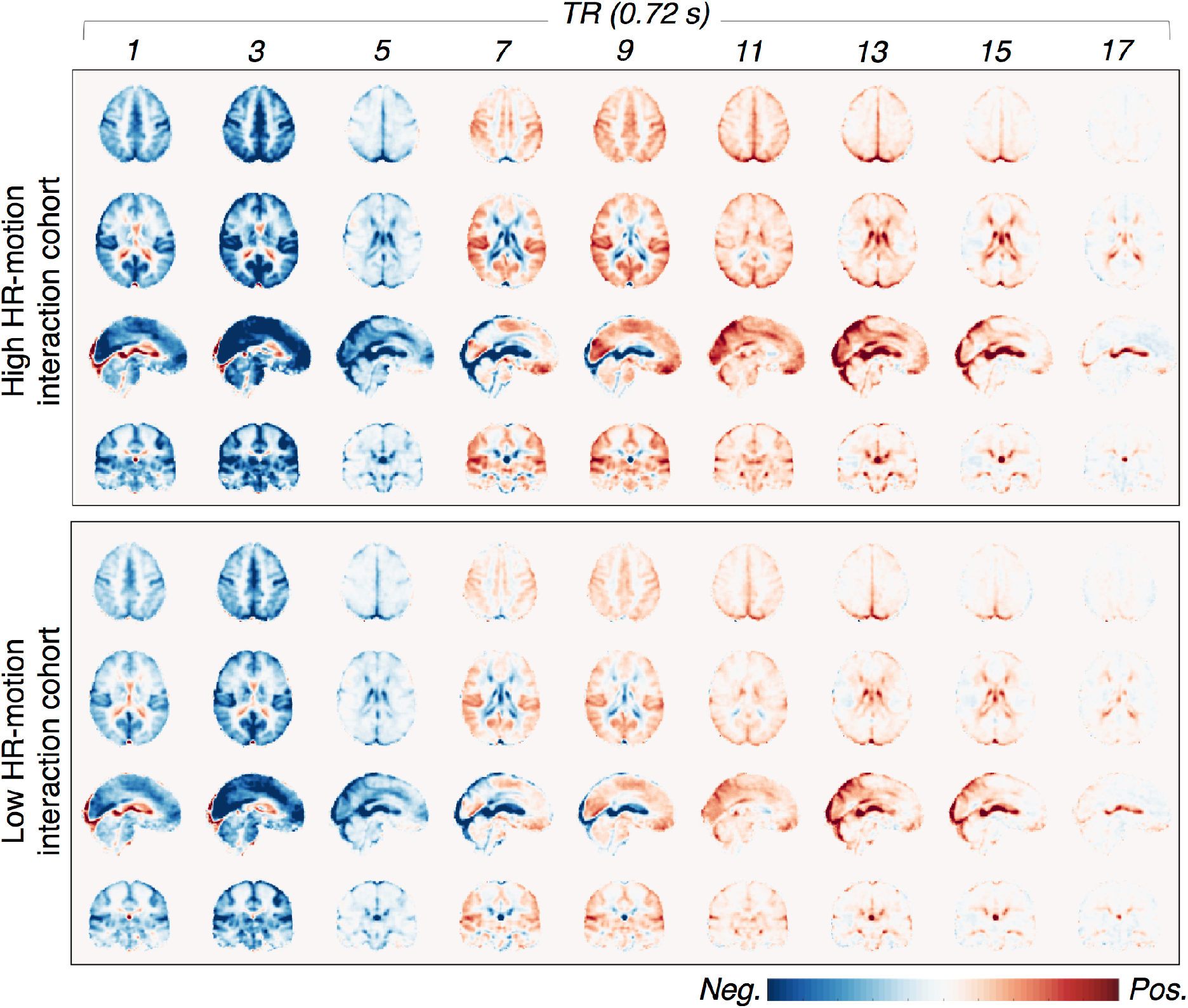
Subjects were separated into two cohorts according to the interaction between head motion and HBI measurements, quantified as the absolute value of the linear Pearson correlation between the RMS of head movements (‘*Movement_AbsoluteRMS.txt*’ released by HCP) and HBI changes, |*r*_HBI-motion_|. iCRFs were averaged within two separate cohorts of subjects with different extents of HBI-motion interactions: ‘high HBI-motion interaction cohort’ includes the 95 subjects with higher |*r*_HBI-motion_| values, mean±stdev = 0.20±0.09; and ‘low HBI-motion interaction cohort’ includes the 95 subjects with lower |*r*_HBI-motion_| values, mean ± stdev is 0.039 ± 0.024. No prominent differences can be observed between these two groups except for changes in response intensities.

**Fig. S8:**
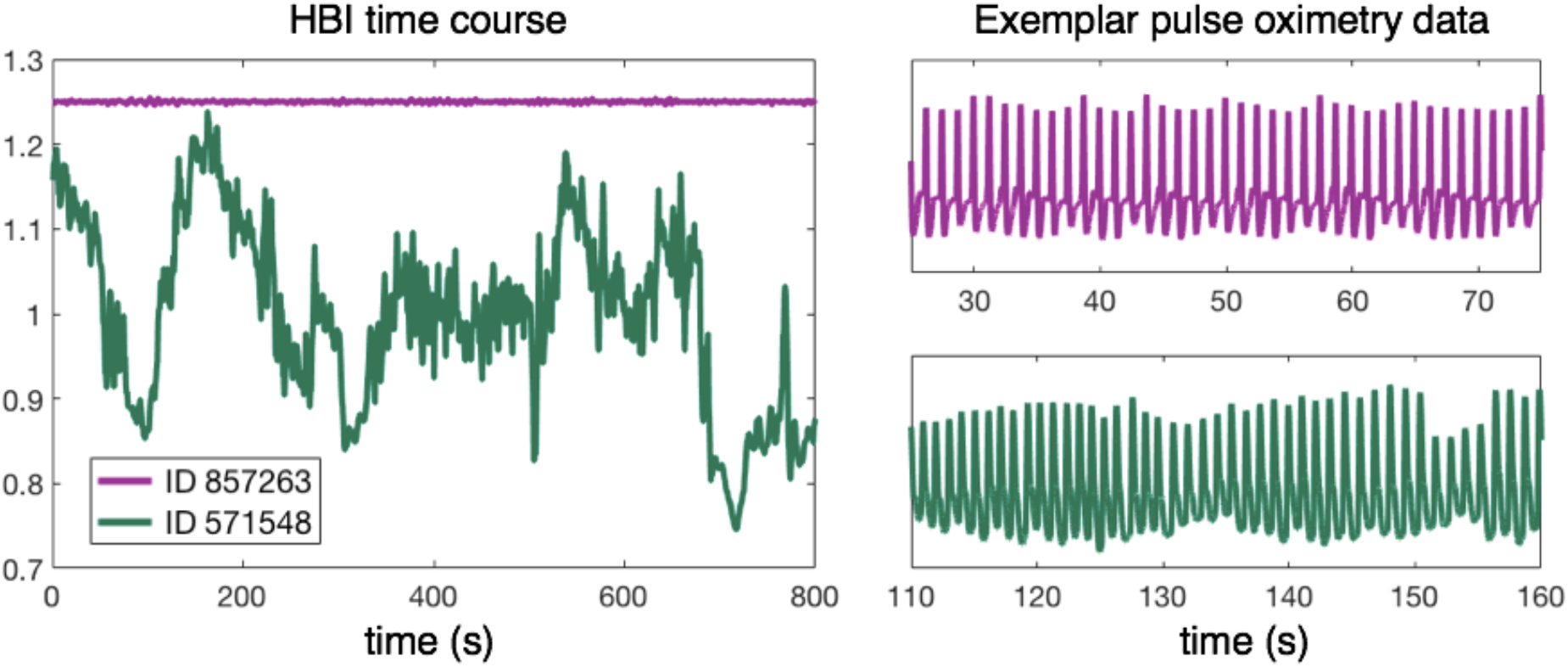
Influence of the level of heart rate variability (HRV) on RRF and iCRF dynamics. HRV can index the healthiness and physiological states of volunteers. We notice that the HCP subjects exhibited a broad range of HRV levels: some subjects had no prominent changes in heart rates (Fig. S7 ID 857263), whereas the others showed notable variability throughout the scan (Fig. S7 ID 571548). Exemplar physiological recordings with distinct levels of HRV are shown. To evaluate the influence of HRV levels on the spatiotemporal patterns of observed physiological dynamics, subjects were separated into two cohorts according to the extents of heart rate variability, quantified as the root mean square of the successive differences (RSSMD) of HBIs. RRFs and iCRFs were averaged within two separate cohorts of subjects with different extents of HBI variability: ‘high HRV cohort’ includes the 95 subjects with higher HRV: mean±stdev of RSSMD is 15.3±7.7 ms; and ‘low HRV cohort’ includes the 95 subjects with lower HRV: mean±stdev of RSSMD is 5.6±2.2 ms. No prominent differences can be observed between these two groups except for notable changes in response intensities.

**Fig. S9:**
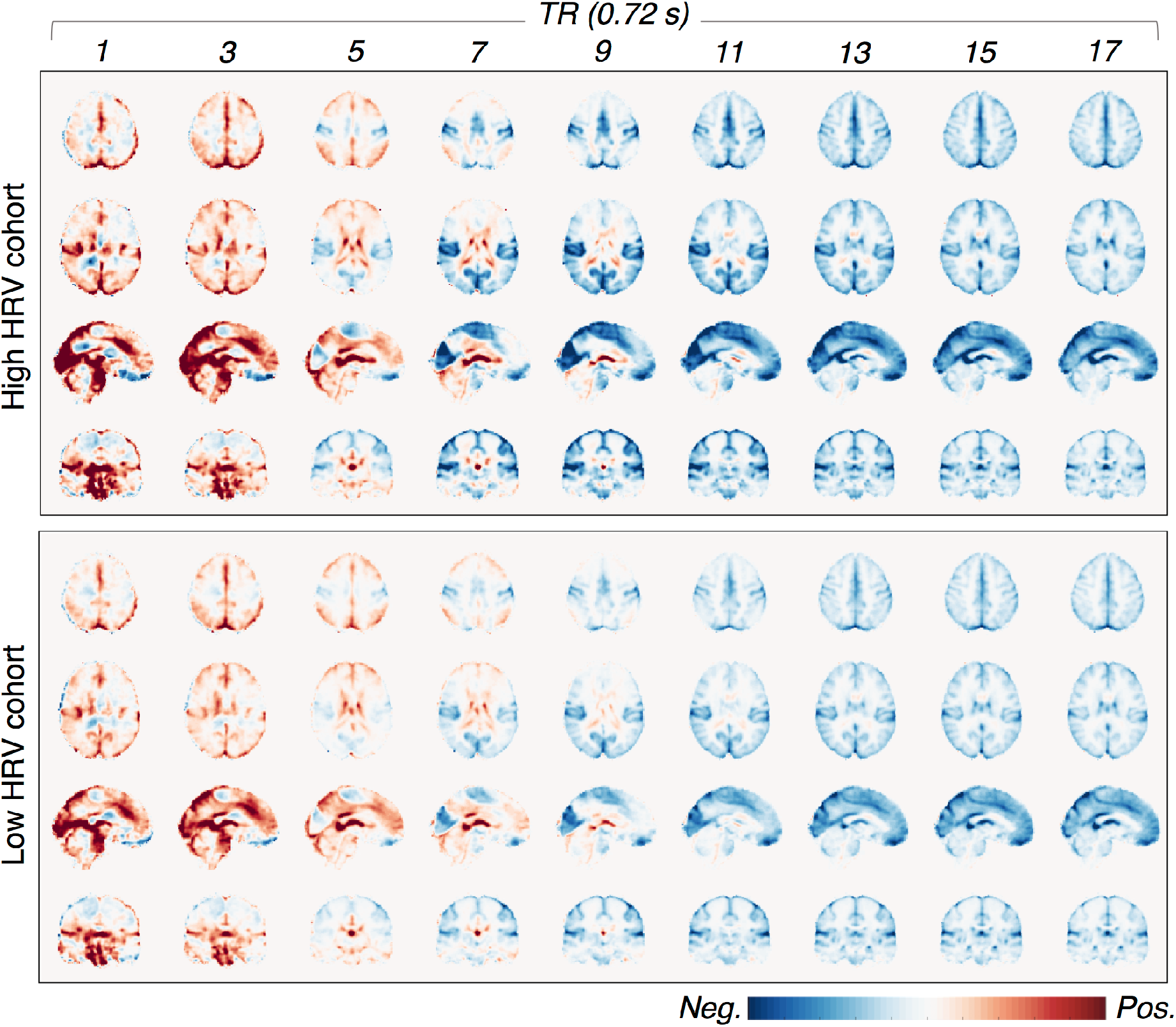
Voxel-wise RRFs averaged within each cohort of subjects based on heart-rate variability, demonstrating similar response functions in both cohorts.

**Fig. S10:**
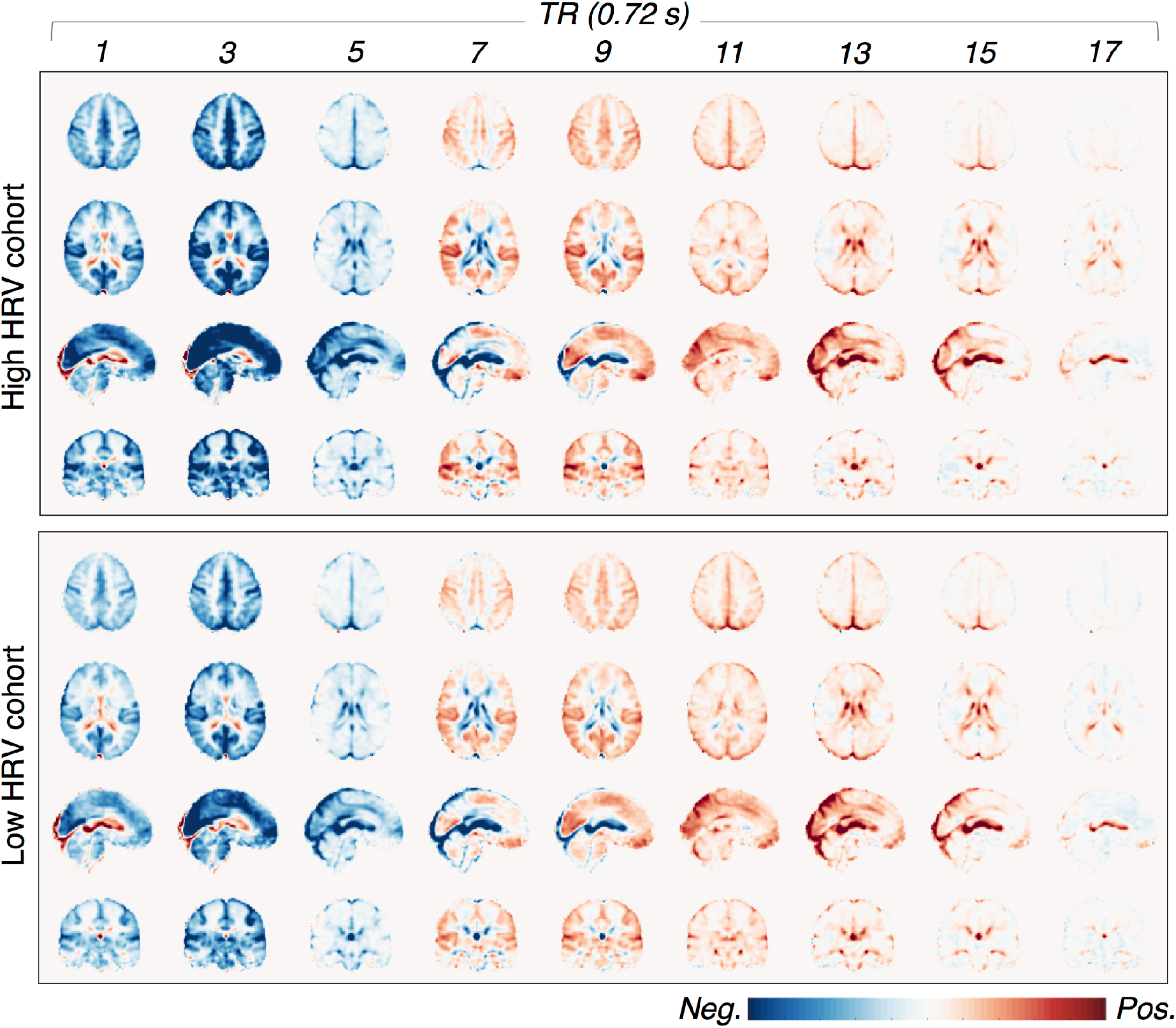
Voxel-wise iCRFs averaged within each cohort of subjects based on heart-rate variability, demonstrating similar response functions in both cohorts.

**Table S1:**
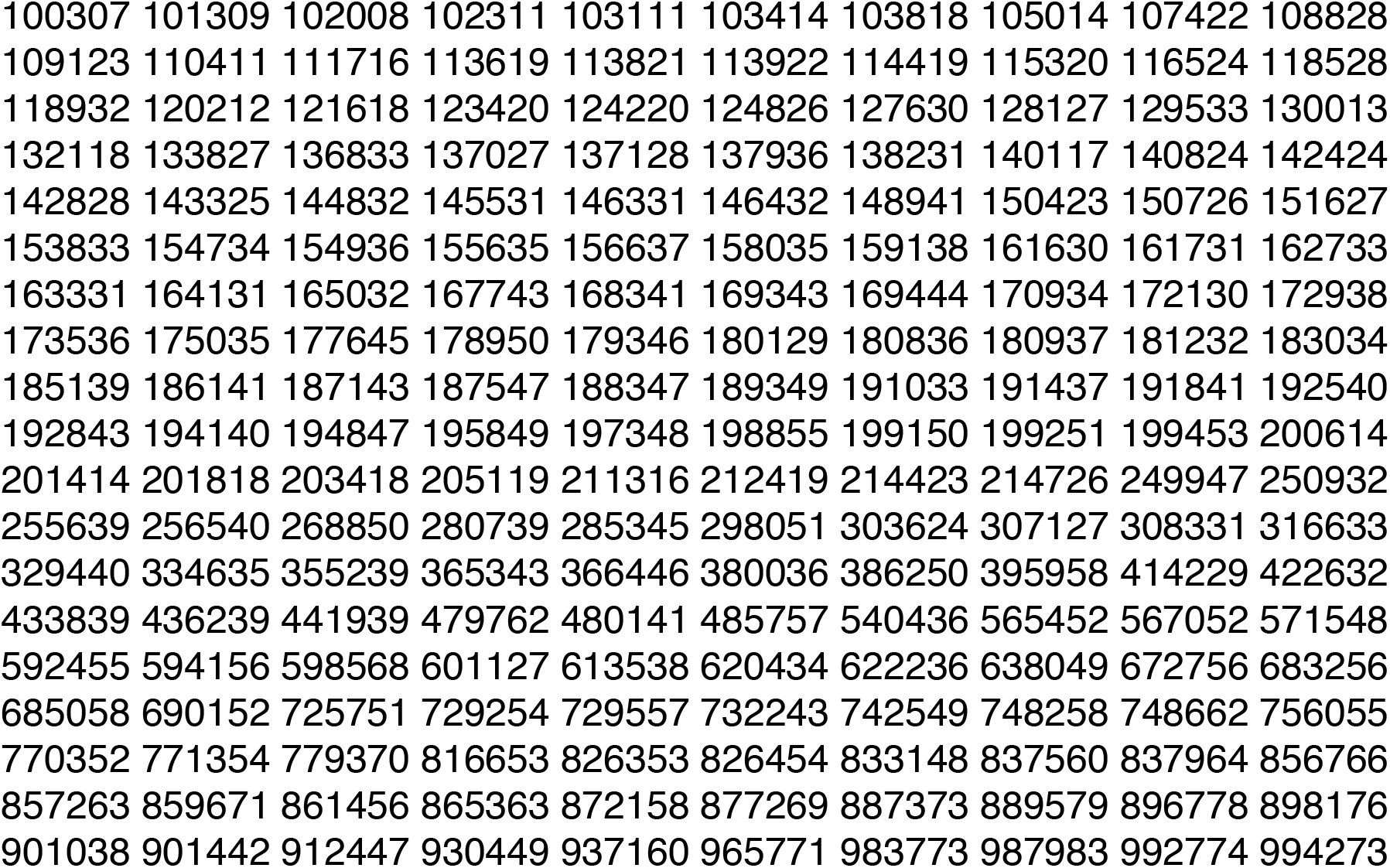
ID numbers of the HCP subjects included in the present study:

**Eqn. S1: analytical forms of the physiological basis functions (Fig. 1C)**

All basis functions were temporally normalized (divided by its maximum value of absolute fluctuations, i.e., max (|*RRF_x_(t) or CRF(t)*|) =1).

RRF basis: *RRF_p_(t)* (primary); *RRF_t_*_1_(*t*),*RRF_t_*_2_(*t*) (temp. deriv.); and *RRF_d_*_1_(*t*),*RRF_d_*_2_(*t*) (disp. deriv.);

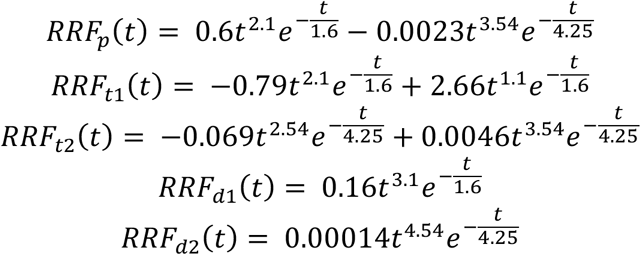

iCRF basis: *CRF_p_*(*t*) (primary); *CRF_t_*_1_(*t*),*CRF_t_*_2_(*t*) (temp. deriv.); and *CRF_d_*_1_(*t*),*CRF_d_*_2_(*t*) (disp. deriv.);

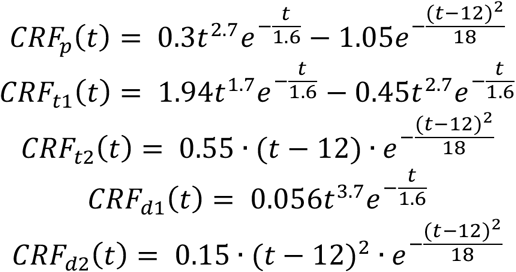

## Footnotes

1 Bright MG, Whittaker JR, Driver ID, Murphy K. 2018. Vascular physiology drives functional brain networks. bioRxiv. doi:10.1101/475491.

2 Kassinopoulos M, Mitsis GD. 2019. Identification of Physiological Response Functions to Correct for Fluctuations in Resting-State fMRI related to Heart Rate and Respiration. bioRxiv. doi: 10.1101/512855.

## Notes

#### Summary of Updates

This version has been revised to update the submission category from "Physiology" to "Neuroscience".

## References

Abbas A, Belloy M, Kashyap A, Billings J, Nezafati M, Schumacher EH, Keilholz S. 2019. Quasi-periodic patterns contribute to functional connectivity in the brain. Neuroimage. 191:193–204.

Aguirre GK, Zarahn E, D’Esposito M. 1998. The variability of human, BOLD hemodynamic responses. Neuroimage. 8:360–369.

Al-Bachari S, Parkes LM, Vidyasagar R, Hanby MF, Tharaken V, Leroi I, Emsley HC. 2014. Arterial spin labelling reveals prolonged arterial arrival time in idiopathic Parkinson’s disease. Neuroimage Clin. 6:1–8.

Alkire MT, Haier RJ, Barker SJ, Shah NK, Wu JC, Kao YJ. 1995. Cerebral metabolism during propofol anesthesia in humans studied with positron emission tomography. Anesthesiology. 82:393–403; discussion 327A.

Alkire MT, Haier RJ, Shah NK, Anderson CT. 1997. Positron emission tomography study of regional cerebral metabolism in humans during isoflurane anesthesia. Anesthesiology. 86:549–557.

Baker AP, Brookes MJ, Rezek IA, Smith SM, Behrens T, Probert Smith PJ, Woolrich M. 2014. Fast transient networks in spontaneous human brain activity. Elife. 3:e01867.

Beissner F, Meissner K, Bar KJ, Napadow V. 2013. The autonomic brain: an activation likelihood estimation meta-analysis for central processing of autonomic function. J Neurosci. 33:10503–10511.

Bernier M, Cunnane SC, Whittingstall K. 2018. The morphology of the human cerebrovascular system. Hum Brain Mapp. 39:4962–4975.

Birn RM, Diamond JB, Smith MA, Bandettini PA. 2006. Separating respiratory-variation-related fluctuations from neuronal-activity-related fluctuations in fMRI. Neuroimage. 31:1536–1548.

Birn RM, Smith MA, Jones TB, Bandettini PA. 2008. The respiration response function: the temporal dynamics of fMRI signal fluctuations related to changes in respiration. Neuroimage. 40:644–654.

Biswal B, Yetkin FZ, Haughton VM, Hyde JS. 1995. Functional connectivity in the motor cortex of resting human brain using echo-planar MRI. Magn Reson Med. 34:537–541.

Black JE, Isaacs KR, Anderson BJ, Alcantara AA, Greenough WT. 1990. Learning causes synaptogenesis, whereas motor activity causes angiogenesis, in cerebellar cortex of adult rats. Proc Natl Acad Sci U S A. 87:5568–5572.

Bonnet MH, Arand DL. 1997. Heart rate variability: sleep stage, time of night, and arousal influences. Electroencephalogr Clin Neurophysiol. 102:390–396.

Boyle PJ, Scott JC, Krentz AJ, Nagy RJ, Comstock E, Hoffman C. 1994. Diminished brain glucose metabolism is a significant determinant for falling rates of systemic glucose utilization during sleep in normal humans. J Clin Invest. 93:529–535.

Bright MG, Murphy K. 2015. Is fMRI “noise” really noise? Resting state nuisance regressors remove variance with network structure. Neuroimage. 114:158–169.

Bright MG, Whittaker JR, Driver ID, Murphy K. 2018. Vascular physiology drives functional brain networks. bioRxiv. doi:10.1101/475491.

Bright MGM, K. 2014. Spatially coupled functional and vascular networks. Proc Intl Soc Mag Reson Med 0441.

Brookes MJ, Hale JR, Zumer JM, Stevenson CM, Francis ST, Barnes GR, Owen JP, Morris PG, Nagarajan SS. 2011. Measuring functional connectivity using MEG: methodology and comparison with fcMRI. Neuroimage. 56:1082–1104.

Buckner RL, Krienen FM, Castellanos A, Diaz JC, Yeo BT. 2011. The organization of the human cerebellum estimated by intrinsic functional connectivity. J Neurophysiol. 106:2322–2345.

Buchsbaum MS, Gillin JC, Wu J, Hazlett E, Sicotte N, Dupont RM, Bunney WE, Jr. 1989. Regional cerebral glucose metabolic rate in human sleep assessed by positron emission tomography. Life Sci. 45:1349–1356.

Byrge L, Kennedy DP. 2018. Identifying and characterizing systematic temporally-lagged BOLD artifacts. Neuroimage. 171:376–392.

Caballero-Gaudes C, Reynolds RC. 2017. Methods for cleaning the BOLD fMRI signal. Neuroimage. 154:128–149.

Cechetto DFSe, C.B. 1990. Role of the cerebral cortex in autonomic function. In: Loewy ADS, K.M., editor. Central Regulation of Autonomic Functions. Oxford: Oxford UP p 208–223.

Chang C, Cunningham JP, Glover GH. 2009. Influence of heart rate on the BOLD signal: the cardiac response function. Neuroimage. 44:857–869.

Chang C, Thomason ME, Glover GH. 2008. Mapping and correction of vascular hemodynamic latency in the BOLD signal. Neuroimage. 43:90–102.

Chang C, O P, de Zwart J, Picchioni D, Chappel-Farley M, Mandelkow H, Duyn, J. 2018. Covariation of pulse oximetry amplitude and BOLD fMRI across vigilance states. Proc Intl Soc Mag Reson Med 0046.

Chen JE, Glover GH, Greicius MD, Chang C. 2017. Dissociated patterns of anti-correlations with dorsal and ventral default-mode networks at rest. Hum Brain Mapp. 38:2454–2465.

Chen JE, Jahanian H, Glover GH. 2017. Nuisance Regression of High-Frequency Functional Magnetic Resonance Imaging Data: Denoising Can Be Noisy. Brain Connect. 7:13–24.

Chen JE, Polimeni JR, Bollmann S, Glover GH. 2019. On the analysis of rapidly sampled fMRI data. Neuroimage. 188:807–820.

Christen T, Jahanian H, Ni WW, Qiu D, Moseley ME, Zaharchuk G. 2015. Noncontrast mapping of arterial delay and functional connectivity using resting-state functional MRI: a study in Moyamoya patients. J Magn Reson Imaging. 41:424–430.

Cordes D, Nandy RR, Schafer S, Wager TD. 2014. Characterization and reduction of cardiac- and respiratory-induced noise as a function of the sampling rate (TR) in fMRI. Neuroimage. 89:314–330.

Critchley HD, Mathias CJ, Josephs O, O’Doherty J, Zanini S, Dewar BK, Cipolotti L, Shallice T, Dolan RJ. 2003. Human cingulate cortex and autonomic control: converging neuroimaging and clinical evidence. Brain. 126:2139–2152.

de la Cruz F, Schumann A, Kohler S, Bar KJ, Wagner G. 2017. Impact of the heart rate on the shape of the cardiac response function. Neuroimage. 162:214–225.

Dripps RD, Severinghaus JW. 1955. General anesthesia and respiration. Physiol Rev. 35:741–777.

Fair DA, Miranda-Dominguez O, Snyder AZ, Perrone AA, Earl EA, Van AN, Koller JM, Feczko E, Klein RL, Mirro AE, Hampton JM, Adeyemo B, Laumann TO, Gratton C, Greene DJ, Schlaggar B, Hagler D, Watts R, Garavan H, Barch DM, Nigg JT, Petersen SE, Dale A, Feldstein-Ewing SW, Nagel BJ, Dosenbach NUF. 2018. Correction of respiratory artifacts in MRI head motion estimates. bioRxiv.

Falahpour M, Refai H, Bodurka J. 2013. Subject specific BOLD fMRI respiratory and cardiac response functions obtained from global signal. Neuroimage. 72:252–264.

Fink BR, Ngai SH, Hanks EC. 1962. The central regulation of respiration during halothane anesthesia. Anesthesiology. 23:200–206.

Finn ES, Shen X, Scheinost D, Rosenberg MD, Huang J, Chun MM, Papademetris X, Constable RT. 2015. Functional connectome fingerprinting: identifying individuals using patterns of brain connectivity. Nat Neurosci. 18:1664–1671.

Fox MD, Raichle ME. 2007. Spontaneous fluctuations in brain activity observed with functional magnetic resonance imaging. Nat Rev Neurosci. 8:700–711.

Fukunaga M, Horovitz SG, van Gelderen P, de Zwart JA, Jansma JM, Ikonomidou VN, Chu R, Deckers RH, Leopold DA, Duyn JH. 2006. Large-amplitude, spatially correlated fluctuations in BOLD fMRI signals during extended rest and early sleep stages. Magn Reson Imaging. 24:979–992.

Glasser MF, Coalson TS, Bijsterbosch JD, Harrison SJ, Harms MP, Anticevic A, Van Essen DC, Smith SM. 2018. Using temporal ICA to selectively remove global noise while preserving global signal in functional MRI data. Neuroimage. 181:692–717.

Glasser MF, Sotiropoulos SN, Wilson JA, Coalson TS, Fischl B, Andersson JL, Xu J, Jbabdi S, Webster M, Polimeni JR, Van Essen DC, Jenkinson M, Consortium WU-MH. 2013. The minimal preprocessing pipelines for the Human Connectome Project. Neuroimage. 80:105–124.

Glover GH, Li TQ, Ress D. 2000. Image-based method for retrospective correction of physiological motion effects in fMRI: RETROICOR. Magn Reson Med. 44:162–167.

Golestani AM, Chang C, Kwinta JB, Khatamian YB, Jean Chen J. 2015. Mapping the end-tidal CO2 response function in the resting-state BOLD fMRI signal: spatial specificity, test-retest reliability and effect of fMRI sampling rate. Neuroimage. 104:266–277.

Golestani AM, Wei LL, Chen JJ. 2016. Quantitative mapping of cerebrovascular reactivity using resting-state BOLD fMRI: Validation in healthy adults. Neuroimage. 138:147–163.

Guyenet PG, Bayliss DA. 2015. Neural Control of Breathing and CO2 Homeostasis. Neuron. 87:946–961.

Handwerker DA, Ollinger JM, D’Esposito M. 2004. Variation of BOLD hemodynamic responses across subjects and brain regions and their effects on statistical analyses. Neuroimage. 21:1639–1651.

He BJ, Snyder AZ, Zempel JM, Smyth MD, Raichle ME. 2008. Electrophysiological correlates of the brain’s intrinsic large-scale functional architecture. Proc Natl Acad Sci U S A. 105:16039–16044.

Hipp JF, Hawellek DJ, Corbetta M, Siegel M, Engel AK. 2012. Large-scale cortical correlation structure of spontaneous oscillatory activity. Nat Neurosci. 15:884–890.

Hudgel DW, Martin RJ, Johnson B, Hill P. 1984. Mechanics of the respiratory system and breathing pattern during sleep in normal humans. J Appl Physiol Respir Environ Exerc Physiol. 56:133–137.

Ikeda K, Kawakami K, Onimaru H, Okada Y, Yokota S, Koshiya N, Oku Y, Iizuka M, Koizumi H. 2017. The respiratory control mechanisms in the brainstem and spinal cord: integrative views of the neuroanatomy and neurophysiology. J Physiol Sci. 67:45–62.

Jahanian H, Christen T, Moseley ME, Pajewski NM, Wright CB, Tamura MK, Zaharchuk G, Group SSR. 2017. Measuring vascular reactivity with resting-state blood oxygenation level-dependent (BOLD) signal fluctuations: A potential alternative to the breath-holding challenge? J Cereb Blood Flow Metab. 37:2526–2538.

Jahanian H, Christen T, Moseley ME, Zaharchuk G. 2018. Erroneous Resting-State fMRI Connectivity Maps Due to Prolonged Arterial Arrival Time and How to Fix Them. Brain Connect. 8:362–370.

Kassinopoulos M, Mitsis GD. 2019. Identification of Physiological Response Functions to Correct for Fluctuations in Resting-State fMRI related to Heart Rate and Respiration. bioRxiv. doi: 10.1101/512855.

Kucyi A, Schrouff J, Bickel S, Foster BL, Shine JM, Parvizi J. 2018. Intracranial Electrophysiology Reveals Reproducible Intrinsic Functional Connectivity within Human Brain Networks. J Neurosci. 38:4230–4242.

Latson TW, McCarroll SM, Mirhej MA, Hyndman VA, Whitten CW, Lipton JM. 1992. Effects of three anesthetic induction techniques on heart rate variability. J Clin Anesth. 4:265–276.

Li Y, Wu P, Lu JT, Chu Y. Hsu Y, Lin F. 2018. Inter-regional BOLD latency after vascular reactivity calibration is correlated to reaction time. Proc Intl Soc Mag Reson Med.0148.

Liu TT. 2016. Noise contributions to the fMRI signal: An overview. Neuroimage. 143:141–151.

Liu TT, Nalci A, Falahpour M. 2017. The global signal in fMRI: Nuisance or Information? Neuroimage. 150:213–229.

Lu H. 2019. Physiological MRI of the brain: Emerging techniques and clinical applications. Neuroimage. 187:1–2.

Lv Y, Margulies DS, Cameron Craddock R, Long X, Winter B, Gierhake D, Endres M, Villringer K, Fiebach J, Villringer A. 2013. Identifying the perfusion deficit in acute stroke with resting-state functional magnetic resonance imaging. Ann Neurol. 73:136–140.

Ma Y, Shaik MA, Kozberg MG, Kim SH, Portes JP, Timerman D, Hillman EM. 2016. Resting-state hemodynamics are spatiotemporally coupled to synchronized and symmetric neural activity in excitatory neurons. Proc Natl Acad Sci U S A. 113:E8463–E8471.

MacIntosh BJ, Filippini N, Chappell MA, Woolrich MW, Mackay CE, Jezzard P. 2010. Assessment of arterial arrival times derived from multiple inversion time pulsed arterial spin labeling MRI. Magn Reson Med. 63:641–647.

MacIntosh BJ, Lindsay AC, Kylintireas I, Kuker W, Gunther M, Robson MD, Kennedy J, Choudhury RP, Jezzard P. 2010. Multiple inflow pulsed arterial spin-labeling reveals delays in the arterial arrival time in minor stroke and transient ischemic attack. AJNR Am J Neuroradiol. 31:1892–1894.

Majeed W, Magnuson M, Hasenkamp W, Schwarb H, Schumacher EH, Barsalou L, Keilholz SD. 2011. Spatiotemporal dynamics of low frequency BOLD fluctuations in rats and humans. Neuroimage. 54:1140–1150.

Mateo C, Knutsen PM, Tsai PS, Shih AY, Kleinfeld D. 2017. Entrainment of Arteriole Vasomotor Fluctuations by Neural Activity Is a Basis of Blood-Oxygenation-Level-Dependent “Resting-State” Connectivity. Neuron. 96:936–948.

Miller KJ, Weaver KE, Ojemann JG. 2009. Direct electrophysiological measurement of human default network areas. Proc Natl Acad Sci U S A. 106:12174–12177.

Mitra A, Snyder AZ, Blazey T, Raichle ME. 2015. Lag threads organize the brain’s intrinsic activity. Proc Natl Acad Sci U S A. 112:E2235–2244.

Murphy K, Birn RM, Handwerker DA, Jones TB, Bandettini PA. 2009. The impact of global signal regression on resting state correlations: are anti-correlated networks introduced? Neuroimage. 44:893–905.

Murphy K, Fox MD. 2017. Towards a consensus regarding global signal regression for resting state functional connectivity MRI. Neuroimage. 154:169–173.

Nofzinger EA, Buysse DJ, Miewald JM, Meltzer CC, Price JC, Sembrat RC, Ombao H, Reynolds CF, Monk TH, Hall M, Kupfer DJ, Moore RY. 2002. Human regional cerebral glucose metabolism during non-rapid eye movement sleep in relation to waking. Brain. 125:1105–1115.

Oken BS, Salinsky MC, Elsas SM. 2006. Vigilance, alertness, or sustained attention: physiological basis and measurement. Clin Neurophysiol. 117:1885–1901.

Ozbay PS, Chang C, Picchioni D, Mandelkow H, Moehlman TM, Chappel-Farley MG, van Gelderen P, de Zwart JA, Duyn JH. 2018. Contribution of systemic vascular effects to fMRI activity in white matter. Neuroimage. 176:541–549.

Pinto J, Nunes S, Bianciardi M, Dias A, Silveira LM, Wald LL, Figueiredo P. 2017. Improved 7 Tesla resting-state fMRI connectivity measurements by cluster-based modeling of respiratory volume and heart rate effects. Neuroimage. 153:262–272.

Power JD, Barnes KA, Snyder AZ, Schlaggar BL, Petersen SE. 2012. Spurious but systematic correlations in functional connectivity MRI networks arise from subject motion. Neuroimage. 59:2142–2154.

Power JD, Plitt M, Gotts SJ, Kundu P, Voon V, Bandettini PA, Martin A. 2018. Ridding fMRI data of motion-related influences: Removal of signals with distinct spatial and physical bases in multiecho data. Proc Natl Acad Sci U S A. 115:E2105–E2114.

Power JD, Plitt M, Laumann TO, Martin A. 2017. Sources and implications of whole-brain fMRI signals in humans. Neuroimage. 146:609–625.

Quaegebeur A, Lange C, Carmeliet P. 2011. The neurovascular link in health and disease: molecular mechanisms and therapeutic implications. Neuron. 71:406–424.

Rangaprakash D, Wu GR, Marinazzo D, Hu X, Deshpande G. 2018. Hemodynamic response function (HRF) variability confounds resting-state fMRI functional connectivity. Magn Reson Med. 80:1697–1713.

Satterthwaite TD, Ciric R, Roalf DR, Davatzikos C, Bassett DS, Wolf DH. 2017. Motion artifact in studies of functional connectivity: Characteristics and mitigation strategies. Hum Brain Mapp.

Shmuel A, Leopold DA. 2008. Neuronal correlates of spontaneous fluctuations in fMRI signals in monkey visual cortex: Implications for functional connectivity at rest. Hum Brain Mapp. 29:751–761.

Shmueli K, van Gelderen P, de Zwart JA, Horovitz SG, Fukunaga M, Jansma JM, Duyn JH. 2007. Low-frequency fluctuations in the cardiac rate as a source of variance in the resting-state fMRI BOLD signal. Neuroimage. 38:306–320.

Smith SM, Beckmann CF, Andersson J, Auerbach EJ, Bijsterbosch J, Douaud G, Duff E, Feinberg DA, Griffanti L, Harms MP, Kelly M, Laumann T, Miller KL, Moeller S, Petersen S, Power J, Salimi-Khorshidi G, Snyder AZ, Vu AT, Woolrich MW, Xu J, Yacoub E, Ugurbil K, Van Essen DC, Glasser MF, Consortium WU-MH. 2013. Resting-state fMRI in the Human Connectome Project. Neuroimage. 80:144–168.

Tak S, Polimeni JR, Wang DJ, Yan L, Chen JJ. 2015. Associations of resting-state fMRI functional connectivity with flow-BOLD coupling and regional vasculature. Brain Connect. 5:137–146.

Tak S, Wang DJ, Polimeni JR, Yan L, Chen JJ. 2014. Dynamic and static contributions of the cerebrovasculature to the resting-state BOLD signal. Neuroimage. 84:672–680.

Tong Y, Frederick BD. 2010. Time lag dependent multimodal processing of concurrent fMRI and near-infrared spectroscopy (NIRS) data suggests a global circulatory origin for low-frequency oscillation signals in human brain. Neuroimage. 53:553–564.

Tong Y, Hocke LM, Fan X, Janes AC, Frederick B. 2015. Can apparent resting state connectivity arise from systemic fluctuations? Front Hum Neurosci. 9:285.

Tong Y, Hocke LM, Nickerson LD, Licata SC, Lindsey KP, Frederick B. 2013. Evaluating the effects of systemic low frequency oscillations measured in the periphery on the independent component analysis results of resting state networks. Neuroimage. 76:202–215.

Tong Y, Lindsey KP, Hocke LM, Vitaliano G, Mintzopoulos D, Frederick BD. 2017. Perfusion information extracted from resting state functional magnetic resonance imaging. J Cereb Blood Flow Metab. 37:564–576.

Triantafyllou C, Hoge RD, Krueger G, Wiggins CJ, Potthast A, Wiggins GC, Wald LL. 2005. Comparison of physiological noise at 1.5 T, 3 T and 7 T and optimization of fMRI acquisition parameters. Neuroimage. 26:243–250.

Triantafyllou C, Polimeni JR, Wald LL. 2011. Physiological noise and signal-to-noise ratio in fMRI with multi-channel array coils. Neuroimage. 55:597–606.

Van de Moortele PF, Pfeuffer J, Glover GH, Ugurbil K, Hu X. 2002. Respiration-induced B0 fluctuations and their spatial distribution in the human brain at 7 Tesla. Magn Reson Med. 47:888–895.

Van Essen DC, Smith SM, Barch DM, Behrens TE, Yacoub E, Ugurbil K, Consortium WU-MH. 2013. The WU-Minn Human Connectome Project: an overview. Neuroimage. 80:62–79.

van Houdt PJ, Ossenblok PP, Boon PA, Leijten FS, Velis DN, Stam CJ, de Munck JC. 2010. Correction for pulse height variability reduces physiological noise in functional MRI when studying spontaneous brain activity. Hum Brain Mapp. 31:311–325.

Vigneau-Roy N, Bernier M, Descoteaux M, Whittingstall K. 2014. Regional variations in vascular density correlate with resting-state and task-evoked blood oxygen level-dependent signal amplitude. Hum Brain Mapp. 35:1906–1920.

Wang J, Rao H, Wetmore GS, Furlan PM, Korczykowski M, Dinges DF, Detre JA. 2005. Perfusion functional MRI reveals cerebral blood flow pattern under psychological stress. Proc Natl Acad Sci U S A. 102:17804–17809.

Webb P. 1974. Periodic breathing during sleep. J Appl Physiol. 37:899–903.

Whittaker J, Driver I, Venzi M, Murphy K. 2018. Blood pressure correlated fluctuations of BOLD origin in fMRI signals: A multi-echo 7T study. Proc Intl Soc Mag Reson Med 0868.

Wise RG, Ide K, Poulin MJ, Tracey I. 2004. Resting fluctuations in arterial carbon dioxide induce significant low frequency variations in BOLD signal. Neuroimage. 21:1652–1664.

Wong CW, Olafsson V, Tal O, Liu TT. 2013. The amplitude of the resting-state fMRI global signal is related to EEG vigilance measures. Neuroimage. 83:983–990.

Yeo BT, Krienen FM, Sepulcre J, Sabuncu MR, Lashkari D, Hollinshead M, Roffman JL, Smoller JW, Zollei L, Polimeni JR, Fischl B, Liu H, Buckner RL. 2011. The organization of the human cerebral cortex estimated by intrinsic functional connectivity. J Neurophysiol. 106:1125–1165.

Yuan H, Zotev V, Phillips R, Bodurka J. 2013. Correlated slow fluctuations in respiration, EEG, and BOLD fMRI. Neuroimage. 79:81–93.

